# A geometric foundation for word meaning in the brain

**DOI:** 10.64898/2026.01.28.702241

**Authors:** Hanlin Zhu, Melissa Franch, Elizabeth A. Mickiewicz, James L. Belanger, Rhiannon L. Cowan, Kalman A. Katlowitz, Ana G. Chavez, Assia Chericoni, Danika Paulo, Xinyuan Yan, Shervin Rahimpour, Ben Shofty, Eleonora Bartoli, Jay A. Hennig, Nicole R. Provenza, Elliot H. Smith, Steven T. Piantadosi, Benjamin Y. Hayden, Sameer A. Sheth

## Abstract

In language models of word meaning, directions in the embedding space often correspond to semantic features that can be reused across different words. For example, a single direction corresponding to gender may differentiate word pairs like “boy/girl”, “uncle/aunt” and “king/queen”. Here we show that the same principle governs semantically driven neural responses in the human brain. We recorded populations of single neurons during podcast listening and identified word sets with consistent meaning differences. Across fifteen sets, including gender, plural, and negation, we observed consistent vectorial directions, resulting in parallelogram structures within the neural manifold. Deviation from parallelism in large language models (LLMs) predicted corresponding deviations in brain-derived parallelism. Among pronouns, vectors corresponding to case, number and person exhibited parallelogram structures individually and, collectively, obeyed the principle of commutativity, resulting in a prismatic structure.

Finally, different semantic variables were preferentially associated with discrete groups of neurons, consistent with energy-efficiency theories. Together, these results establish a geometric foundation for the neural encoding of word meaning.

## INTRODUCTION

Word meanings can be captured by vectorial embeddings that convey their important features (Blei, Ng, & Jordan, 2001; Gärdenfors, 2000; Landauer & Dumais, 1997; Turney & Pantel, 2010). In word models and large language models, these vectorial embeddings are systematically related such that stable features of meaning fall along consistent axes, resulting in parallelogram structures for analogy sets (Mikolov et al., 2013; Pennington et al., 2014). Thus, for example, a *gender* vector that spans the embeddings for the words “*brother*/*sister*” would also span embeddings of “*prince*/*princess*” and “*waiter/waitress*”. As a result, directional vectors can be used constructively to build new meanings and make inferences about the meaning of previously unseen words. This feature is valuable for many aspects of reasoning such as analogical thought, generalization, comparison, and even creativity (Gärdenfors, 2014; Gentner, 1983, 2010; Hofstadter, 1995, 2001; Holyoak et al., 2001; Lake et al., 2017; Piantadosi et al., 2024; Ward et al., 1999).

We currently lack a comprehensive theory for how semantic meaning is represented in the brain (Binder et al., 2009; Patterson et al., 2007). Here we propose that the brain, like LLMs, makes use of axes with interpretable meanings that are used consistently across a range of words. Furthermore, we hypothesize that the variation in the alignment of specific word pairs to a given semantic axis will correspond to analogous variation in LLMs, because both reflect subtle variation in semantic geometry.

We base these hypotheses on growing evidence that semantic representations in the brain share structural features with Large Language Models (LLMs), including high-dimensional distributed embeddings and attention-head structures (Caucheteux & King, 2022; Franch et al., 2025; Katlowitz et al., 2025; Schrimpf et al., 2021). Additionally, LLMs closely mirror human performance on tasks involving semantic directions such as analogical reasoning (Webb et al., 2023; Musker et al., 2025; Grand et al., 2022).

Independent of these hypothesized computational parallels with LLMs, a small set of neuroimaging studies supports the possibility that the cerebral cortex uses stable semantic axes (Zhang et al., 2020; Wu et al., 2022). However, fMRI provides limited insight into neural coding principles for semantics because of its limited spatial and temporal resolution.

A growing body of evidence suggests that neural representations may be organized in a *factorized* (sometimes called *disentangled*) manner, in which distinct latent variables are encoded along independent representational axes (Courellis et al., 2024; Fusi et al., 2016; Rigotti et al., 2013; Bernardi et al., 2020). Such factorized representations enable flexible recombination of representational components, robust cross-condition generalization, and the compositional structure required for abstract reasoning and symbolic-like operations (Fodor & Pylyshyn, 1988; Smolensky, 1990; Fusi et al., 2016; Higgins et al., 2018). One strong test of factorized representations is a *prismatic* organization of neural state space, in which combinations of factors occupy vertices of parallel subspaces whose translations correspond to operations applied along independent dimensions (Trager et al., 2024; Yang et al., 2019; Ostrow & Fiete, 2024). This geometry can in turn give rise to commutativity: applying two transformations in different orders yields equivalent representational displacements (Quessard et al., 2020; Higgins et al., 2018), reflecting the additive structure of the underlying latent variables. Compared to parallelogram structures, prismatic geometry implies a stronger organizational principle: that multiple variables are simultaneously represented in separable subspaces whose interactions obey consistent algebraic rules across the full representational manifold—supporting systematic recombination and generalization (Lake et al., 2017; Baroni, 2020).

To test these ideas, we examined neural encoding of analogical relationships during natural speech listening. Relying on natural speech is essential to determine whether these geometric principles are intrinsic, rather than scaffolded by specialized laboratory tasks currently prevalent in the factorization literature. Importantly, continuous speech captures words in their natural, contextualized state and inherently drives dynamic cognitive processes—such as the spontaneous reactivation of antecedent concepts by pronouns (Dijksterhuis et al., 2024)—allowing us to investigate how natural comprehension shapes underlying geometric structures. The hippocampus is especially interesting in this regard due to its putative role in using geometric principles to embed conceptual representation (Behrens et al., 2018; Bernardi et al., 2020; Constantinescu et al., 2016; Courellis et al., 2024; Kafkas et al., 2024; Mack et al., 2018; Theves et al., 2020), including for language semantics (Blank et al., 2016; Davachi, 2006; Wolna et al., 2025). We and others have argued that its role includes encoding word meanings during speech listening (Franch et al., 2025; Katlowitz et al., 2025; Dijksterhuis et al., 2024; Blank et al., 2016). Indeed, we previously showed that *distances* in neural coding space correspond to semantic distance; here we ask whether *directions* may have semantic meanings. However, the hippocampus is not the only region associated with semantic coding (Huth et al., 2016; Tang et al., 2023) or geometric abstraction (Bernardi et al., 2020); we therefore further explored the generality of these effects by examining responses in two other brain regions, the anterior cingulate cortex (ACC) and orbitofrontal cortex (OFC) and identified regional specialization of semantic directions.

## RESULTS

Fourteen native English-speaking patients (6 males and 8 females) undergoing neural monitoring for epilepsy listened to 47 minutes of English speech (six monologues taken from the Moth podcast, **Methods**, **Figure 1A** and **B**, Franch et al., 2025). We collected responses of isolated single neurons in the hippocampus (HPC, n = 437 neurons, 14 patients), the anterior cingulate cortex (ACC, n = 241 neurons, 12 patients), and the orbitofrontal cortex (OFC, n = 51 neurons, 5 patients). Many neurons showed a clear response to the onset of individual words, with a characteristic ramping to a peak, followed by a gradual decline in firing (**Figure 1C**, **Supplementary Figure 1**, Franch et al., 2025). We computed firing rates during a time window starting 80 ms after the onset of each word (to account for the approximate response latency) and lasting the duration of the word plus 40 ms.

**Figure 1.**
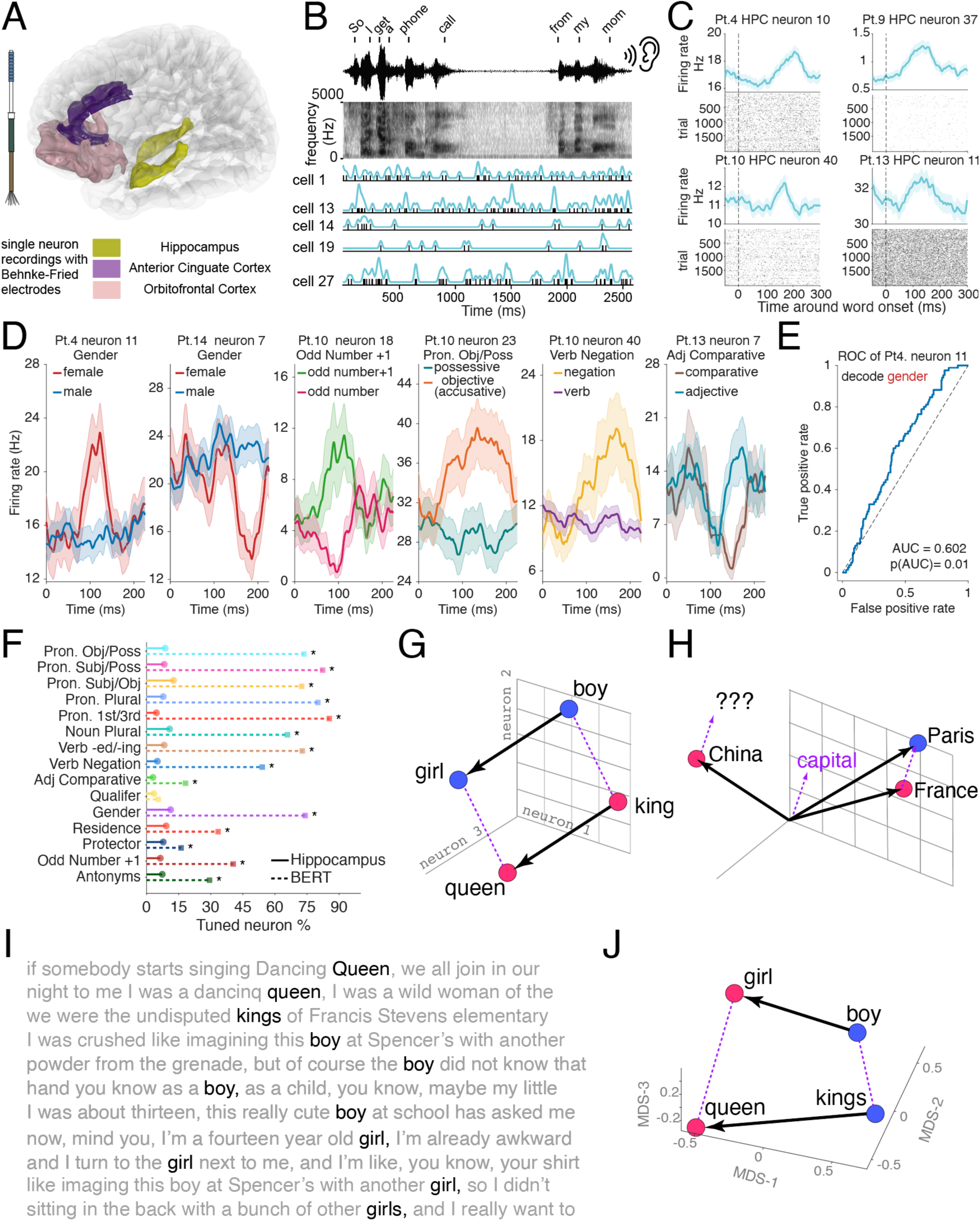
| Hippocampal single neurons encode semantic axes and support analogical vector geometry during the perception of natural speech. **A,** Recording setup and scale. 14 epilepsy patients were implanted with Behnke–Fried depth electrodes. We recorded single units from HPC, ACC and OFC. **B,** Experiment schematic showing continuous speech segment (waveform and spectrogram) with aligned spiking activity from five simultaneously recorded hippocampal neurons. Tick marks denote spike times; blue traces show smoothed firing rate. Word boundaries/onsets are indicated above the waveform. **A** and **B** partially adapted from Franch et al., 2025 with permission. **C,** Peri-word onset responses for four example hippocampal neurons (Pt. = patient). Top: trial-averaged firing rate aligned to word onset (dashed line at 0 ms; shading indicates ± s.e.m.). Bottom: raster plots across word presentations **D,** Example hippocampal neurons whose firing rates differentiate word pairs along specific semantic axes identified in the stimulus text. Left to right: gender (female vs male; Pt.4 neuron 11 and Pt.14 neuron 7), odd number vs odd number+1 (Pt.10 neuron 18), pronoun objective vs possessive (Pt.10 neuron 23), verb negation vs affirmative verb forms (Pt.10 neuron 40), and comparative vs positive (base) adjectives (Pt.13 neuron 7). Traces show mean firing rate aligned to word onset with shading (± s.e.m.); n(male) = 207 trials, n(female) = 85, n(odd number) = 45, n(odd number +1) = 34, n(objective) = 151, n(possessive) = 246, n(verb) = 384, n(negation) = 61, n(adjective) = 21, n(comparative) = 14 trials. **E,** Receiver operating characteristic (ROC) analysis for decoding gender from the firing rate of Pt.4 neuron 11. Significance computed relative to a label-shuffled null distribution (1000 permutations). **F,** Proportion of hippocampal neurons tuned to each of 15 semantic relationships (solid segments) compared to the proportion of tuned units (see Results for definition) in the last layer of BERT (dotted segments; 768 dimensions). Tuning was assessed using the same AUC-based method applied to neurons and model units (multiple-comparisons correction across 15 analogies). Asterisks denote categories with significant differences between hippocampus and BERT (two-proportion Z-test, q < 0.05; correction = FDR). **G,** Schematic illustrating parallelogram geometry expected when a semantic axis is represented consistently: the displacement vector *“boy”*→*“girl”* parallels *“king”*→*“queen”* in an embedding space. **H,** Schematic of analogical vector arithmetic (e.g., *“Paris”* − *“France”* + *“China”* ≈ ?/*“Beijing”*), illustrating how consistent axes enable analogy completion in a novel condition. **I,** Example stimulus excerpt showing repeated occurrences of target lemma (black) across distinct contexts (grey). **J,** Neural embeddings of *“boy”*, *“girl”*, *“kings”*, and *“queen”* computed from the population responses of 100 gender-selective neurons and visualized using multidimensional scaling. Translation vectors show that *“kings”*→*“queen”* aligns with *“boy”*→*“girl”*.

### Hippocampal neural populations have aligned semantic axes

We systematically searched the podcast text (7,346 total words; 1,351 unique words) to identify word pairs that differed along a single semantic direction, or axis. We identified fifteen such semantic directions, which applied to 206 word pairs (summarized in **Table 1**; full list in **Supplementary Table 1**). We define these semantic directions as “analogical relationships,” and we use the terms interchangeably. These analogical relationships were identified before inspection of the data and were based on well-known analogical categories from the literature. They included both semantic (e.g., gender) and grammatical (e.g., case) analogies (**Table 1**).

**Table 1.**
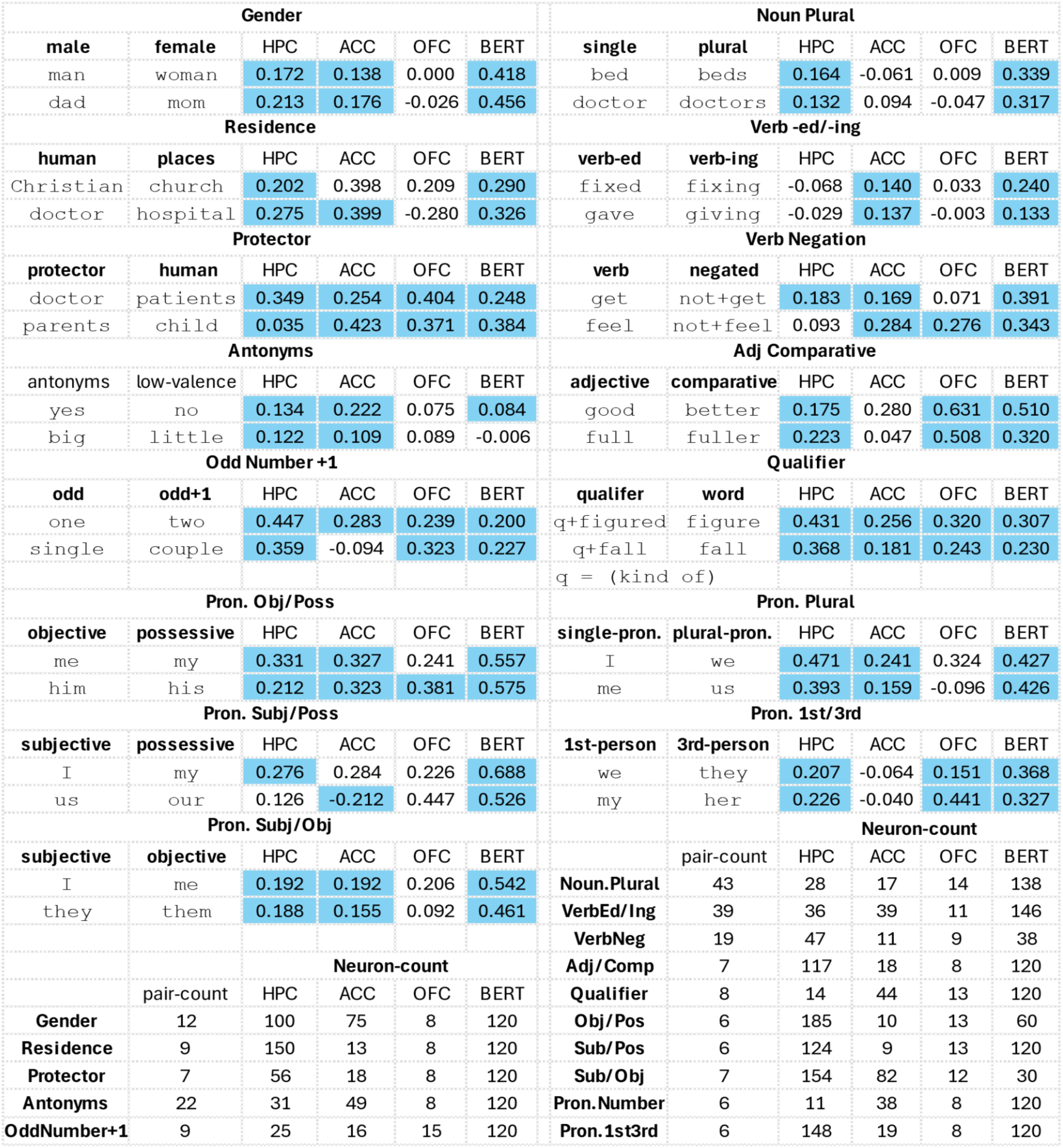
Example word pairs and pair/neuron count summary. Top: Example word pairs of each analogy and their mean cosine similarities (after subtraction) with the other word pairs in the same analogy across different brain areas/LLM. Light blue means significantly aligned pairs. Bottom: number of word pairs and the number of neurons with strong tuning strength (low p(AUC) value) used for each analogy.

We focus first on hippocampal neurons given their putative role in conceptual representations. We found individual hippocampal neurons whose responses systematically differed between the two classes within an analogy (**Figure 1D**). For example, neuron 4.11 (i.e., patient 4, neuron 11) exhibited larger responses to twelve female-associated terms (e.g., “*girl*”, “*queen*”) than to the corresponding twelve male terms (“*boy*”, “*king*”). Other neurons distinguished quantitative or grammatical modifiers. Neuron 10.18 differentiated between odd numbers and their immediate even successors (n and n+1, e.g., “*three*” vs. “*four*”), while neuron 13.7 differentiated comparative forms of adjectives from their stem forms (technically, the positive form (e.g., “*cooler*” vs. “*cool*”)). Meanwhile, neuron 10.40 differentiated negated from non-negated (i.e., affirmative) verbs (e.g., “*not come*” vs. “*come*”), and neuron 10.20 separated possessive from accusative pronouns (e.g., “his” vs. “him”). These example neurons raise the possibility that the brain may make use of consistent response axes for semantic information.

We quantified the firing rate separation between these groups at individual neuron level using a balanced AUC (area under the curve) metric (**Figure 1E**, **Methods**, Franch et al., 2025). A neuron’s tuning strength was defined with its AUC p-value against a label-shuffled null distribution. In this example (**Figure 1E**), neuron 4.11 exhibited a significant (p = 0.01) but modest AUC of 0.6 (chance = 0.5), indicating above chance coding power to differentiate between male and female. This method is agnostic about which of the two conditions elicits a higher firing rate. For comparison, we quantified the proportion of modulated units in the last layer of BERT, a large language model (LLM) that serves as a benchmark for statistical learning of semantic relationships (Devlin et al., 2019; Rogers et al., 2020; Schrimpf et al., 2021, **Figure 1F**). Concretely, the BERT model we analyzed is a stack of transformer encoder layers that outputs, for each input word at each layer, a 768-dimensional contextual hidden state (embeddings); we treat each of the 768 dimensions as an artificial “unit” whose activation for that word is analogous to an instantaneous firing rate. We found that BERT generally possesses a higher density of units aligned with these semantic axes (two-sided two-proportion Z-test, FDR corrected, ɑ = 0.05, q < 0.05). Specifically, it has a greater number of tuned units in 14 out of the 15 analogy categories (all but qualifiers) tested when using the same words in the same contexts (See **Table 1** for complete examples and words used in each of the 15 analogies). Thus, the human hippocampus (7.62% ± 0.72% tuned neurons, mean ± s.e.m., n = 15 categories) more sparsely encodes these semantic axes compared to an LLM (53.59% ± 7.03%). Notably, however, layer-wise scans of BERT and GPT-2 (**Supplementary Figure 2**) reveal that the fraction of units meeting our analogy-tuning criterion generally decreases with increasing layer depth. This may suggest a progressive compression of axis-aligned information into fewer internal units, pointing toward a possible convergence with the sparse coding regime of the brain in deeper model layers.

If ensemble responses to words like “*boy*,” “*girl*,” “*king*,” and “*queen*” are vectors whose features are coded as vectors, then this quartet of related words will have a parallelogram shape on the neural manifold (**Figure 1G**, Rumelhart & Abrahamson, 1973; Mikolov et al., 2013; Allen & Hospedales, 2019; Peterson et al., 2020). Parallelogram-shaped representations have many convenient properties; for example, they allow vector arithmetic to solve analogical reasoning problems (**Figure 1H**, Rumelhart & Abrahamson, 1973).

In our stimulus set, the words “*boy*” and “*girl*” were each repeated four times, in various contexts (**Figure 1I**). We calculated the average response evoked by these words from our a priori gender-selective neurons (that is, neurons with high gender tuning strength, defined as p(AUC) < 0.125, n = 100 out of 437 neurons), and defined the results as the ***neural embeddings*** of the two words, respectively. We then used the same procedure to derive neural embeddings for “*kings*” and “*queen*” using the same set of neurons.

We found that the direction of the *king→queen* vector is more similar to the average *boy→girl* vector (cosine similarity = 0.57) than to a random word*→*random word vector (cosine similarity = 0.007 ± 0.003, mean ± s.e.m., n = 10000 random word pairs, p = 0.013, permutation test, **Supplementary Figure 3**). To see this result, we plot the vectors, using multidimensional scaling (MDS, Carroll & Arabie, 1980; see **Methods**) to facilitate visualization (**Figure 1J**).

### Gender-based word pairs have an aligned axis

We next extended the analysis to the eleven other word pairs having the same gender relationship as “*boy*”/“*girl*.” Visual inspection shows that gender pairs are mostly aligned along a single axis (**Figure 2A and B**). We quantified this alignment using the high-dimension neural population difference vectors. (Note that we used full high dimension vectors, rather than MDS-reduced ones shown in the figures to avoid any risk that the MDS procedure artifactually aligns vectors). To do this, we systematically removed each word pair and calculated the average angular distance (cosine similarity) between that pair and all others. To determine significance, we calculated a null distribution from random word differences and applied the Benjamini-Hochberg false discovery rate (FDR) correction for multiple word pairs tested. (Benjamini & Hochberg, 1995).

**Figure 2.**
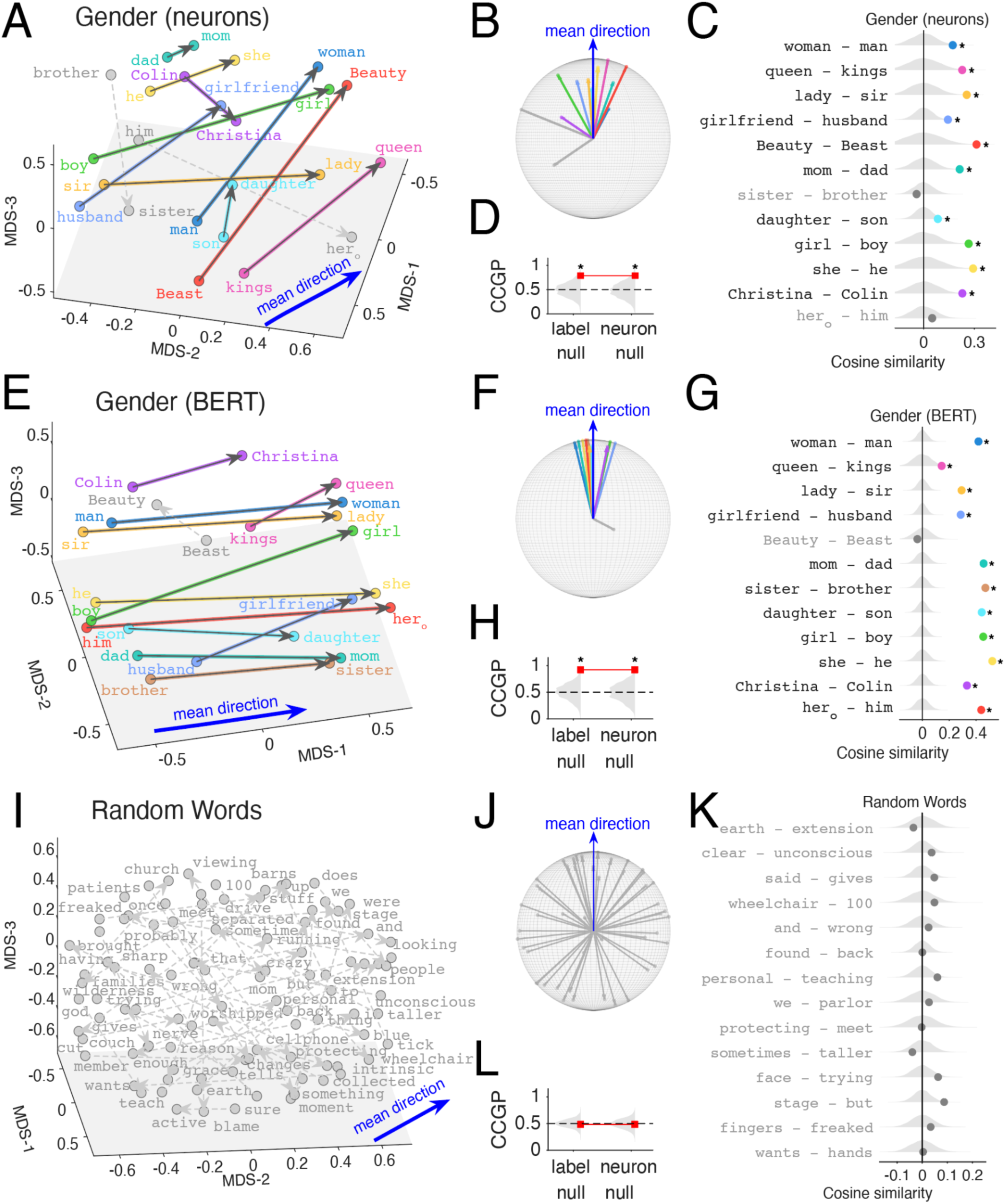
| Word pairs sharing a semantic relationship form aligned directions in hippocampal population space and in an LLM. **A,** MDS visualization of neural embeddings for the gender analogy set (12 female–male word pairs). Each point is a word, and each arrow is the difference vector oriented from the male-associated term to the female-associated term. Colors identify individual word pairs and are used consistently across **A–C**. Grey points/arrows indicate pairs that do not reach significance in the alignment test in **C**. The grey plane is a visual aid for depth only (no interpretive meaning). Hero means the objective form of “*her*” rather than the possessive form. **B,** A visualization of the same vectors in **A** after translating all vector tails to a common origin (tail alignment), applying a rigid rotation so the mean direction (blue) points upward, and normalizing vector lengths to the unit sphere (visualization only; not a separate analysis). Colors correspond to the same pairs shown in **A** and **C**. **C,** Quantification of vector alignment for each gender pair using cosine similarity computed in the high dimensional neural population space (not the MDS-reduced space). For each pair, the dot shows the leave-one-out mean cosine similarity between that pair’s high-dimensional difference vector and the remaining gender vectors; dot colors correspond to the pair colors in **A–B**. Grey violins indicate the null distribution derived from random word-pair differences (n = 12 random pairs, 10000 draws). Asterisks denote significance after Benjamini–Hochberg FDR correction across the 12 tested pairs (q < 0.05). Pair labels are written as female − male. **D,** Cross-condition generalization performance (CCGP) for gender in the neural population. A linear classifier is trained to discriminate gender using trials from a subset of word pairs and tested on a held-out pair (leave-one-pair-out; observed accuracy = 0.788, red markers/line). Grey violins show two null distributions: a label-shuffle null (shuffling gender labels; n = 600 shuffles) and a neuron-shuffle null (shuffling neuron identities while preserving per-neuron firing statistics; n = 600 shuffles). The dashed line indicates chance (0.5). p(label shuffle) = 0.022; p(neuron shuffle) = 0.032. **E–H,** Same analyses as **A–D**, applied to contextual token embeddings from BERT using the same word tokens/contexts. **E,** MDS visualization of BERT embeddings for the gender analogy set (colors correspond to the same word pairs as in **A**; grey indicates the pair(s) not significant in **G**). **F,** Tail-aligned and unit-normalized BERT difference vectors, rotated so the mean direction points upward (visualization only). **G,** Leave-one-out cosine similarity alignment test in BERT, shown with the same conventions as **C** **H,** CCGP for gender same as in D but for BERT embedding space compared against label-shuffle and unit-shuffle null distributions **I–L,** Control analyses on 50 random word pairs. **I,** MDS visualization of random word neural embeddings in HPC and their pairwise difference vectors (grey plane shown for depth cue only). **J,** Tail-aligned and unit-normalized random difference vectors exhibit broad angular dispersion on the unit sphere. **K,** Leave-one-out cosine similarity values for example random pairs compared to the null distribution (grey violins), showing no systematic alignment (stars absent after FDR; q > 0.05). 15 example words shown, see Supplementary Figure 3 for the full list. **L,** CCGP for random pairs near chance and not exceeding null distributions.

We found that neural difference vectors for valid analogical pairs were more aligned with the word pairs in the same analogy category (**Figure 2C**) than random pairs were. The significant pairs included “*mom*”/“*dad*,” “*daughter*”/“*son*,” and “*Beauty*”/“*Beast*” (referring to characters in the Disney movie), as well as the approximate pair “*girlfriend*”/“*husband*”. Two of the twelve pairs (“*sister*”/“*brother*,” q = 0.77 and “*her*”/“*him*,” q = 0.26) did not reach statistical significance. However, the overall proportion of *a priori* identified pairs (n = 10/12) was much greater than chance (p < 0.0001, two-sided binomial test).

This pattern is observed even if we restrict our comparison set to all nouns (p < 0.0001), indicating that our result is not an artifact of the part of speech of the control set having a single shared axis.

To assess the robustness of our axis-alignment results against different similarity definitions (which we currently use cosine similarity) and evaluation regime, we recomputed alignment using Euclidean and Chebyshev similarity and additionally tested an analogy-retrieval paradigm (ranking the target under B−A+C); none of these changes altered our overall conclusions on the number of significant aligned pairs across different analogies. (**Supplementary Figure 4**; Wilcoxon signed-rank tests, q(cosine, Euclidean) = 0.07; q(cosine, Chebyshev) = 0.22; p(cosine, retrieval) = 0.097).

The parallel organization of semantic axes for gender supports generalization (**Figure 2A-C**); however parallelism alone does not guarantee cross-condition generalization, since generalization can be limited by the relative positioning of training vs. test conditions (e.g. test cases being packed closer than the training cases) and by trial-to-trial variability (Bernardi et al., 2020). We next assessed the cross condition generalization performance (CCGP) for neural responses across categories (Bernardi et al., 2020; see also Tang et al., 2023; Courellis et al., 2024). Unlike leave-one-out decoding, cross-condition generalization performance tests whether a neural representation supports systematic generalization across words, thereby assessing abstract, factorized population codes. In short, CCGP quantifies the ability of a linear readout to recover an abstract relationship (e.g., gender) from a novel word pair it has never seen before. To do so, we trained a linear classifier to distinguish between opposing poles of an analogy (e.g., male/female) using a subset of word pairs (e.g., “*king*”/“*queen,*” “*boy*”/“*girl*”) and then tested its ability to correctly classify individual trials from a completely held-out pair (e.g., “*man*”/“*woman*”). CCGP requires that the neural decision boundary learned from specific examples must transfer to novel word pairs. We found that the hippocampal population code supports significant cross-condition generalization for gender (**Figure 2D**). The classifier achieved high accuracy (0.788) on held-out pairs (red line). Crucially, this performance exceeded not only the standard chance level derived from shuffling class labels "label null" (p(label shuffle) = 0.022), but also a more rigorous "neuron-shuffle" null (p(neuron shuffle) = 0.032), which preserves the firing rate statistics of individual neurons while destroying their consistent identity across words, supporting an abstract geometry of gender (Courellis et al., 2024).

These patterns are similar to, but less strong, than the ones observed in a large language model using the same stimulus set (BERT, Devlin et al., 2019). Specifically, BERT vectors were very tightly aligned (**Figure 2E-G**), with the exception of “*Beauty*”/“*Beast*”, presumably due to their rarity in the training set. The lower performance of the neural data relative to the LLM is consistent with the high noise observed in neural responses, as well as the fact that the hippocampus is undoubtedly involved in a variety of other tasks aside from language processing.

Finally, as a control, we repeated the same analysis on a set of 50 randomly chosen word pairs, randomly split in half and labelled each half as either “male” or “female” to validate that we don’t see alignment for just any set of words and randomly defined tuned neurons. We do not (**Figure 2I-K**). For example, vectors calculated based on randomly chosen words “*earth*”/“*extension*” don’t predict responses to “*clear*”/“*unconscious*” or “*said*”/“*gives*”. Indeed, the randomly chosen word pairs show a broad diversity of vector angles, consistent with their random selection.

### Semantic axes generalize to diverse semantic relations

We next tested six other semantic relationships (**Figure 3A-C**). For example, we *a priori* identified ten word pairs reflecting the abstract semantic relationship that we call **residence**, corresponding to a place and a person or group that is associated in any way with that place. These included “*doctor*”/“*hospital*,” “*Texans*”/“*Texas*,” and “*Christian*”/“*church*.” Among the ten pairs in our stimulus set, all were significantly more aligned than chance (q < 0.05, permutation test), except for “*family*”/“*home*” (q = 0.12). Overall, the proportion of aligned pairs was much greater than chance (p < 0.0001, binomial test; CCGP: p < 0.05).

**Figure 3.**
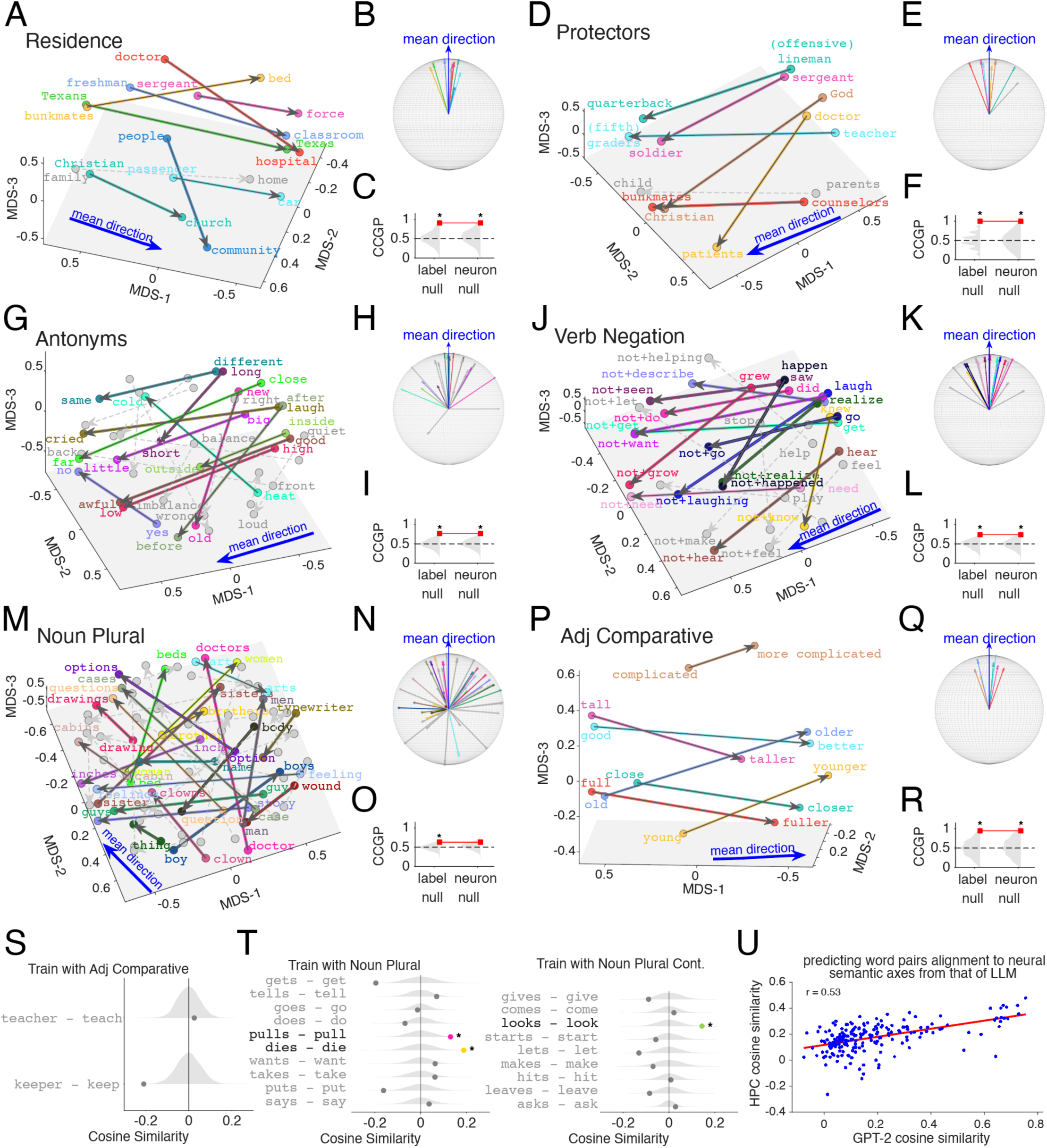
**| Diverse analogical relations are structured along distinct semantic axes** **A–R**, Six semantic relationships define consistent difference directions in hippocampal population space and support cross-condition generalization. For each relationship, the left panel shows a 3D MDS visualization of neural word embeddings with arrows indicating paired difference vectors (direction defined by the relationship; MDS is for visualization only). Colored arrows/labels denote pairs whose high-dimensional difference vectors are significantly aligned with other pairs in the same analogy (permutation test against a random word-pair null; Benjamini–Hochberg FDR across pairs within category); grey arrows/labels denote non-significant pairs. The blue arrow shows the mean direction projected into the MDS view. Grey planes are visual aids only (no interpretive meaning). Middle panels show the same vectors after tail alignment to a common origin and rigid rotation, so the mean direction points upward, with vector lengths normalized to a unit sphere (visualization only; angles are preserved). Right panels show cross-condition generalization performance (CCGP) for each relationship (red marker), compared with a label-shuffle null and a neuron-shuffle null (grey violins; dashed line indicates chance). Asterisks indicate p < 0.05 (permutation test; n shuffles = 600) relative to the corresponding null. See Supplementary Figure 3 for a full list of word-pair labels in the MDS plot, color-coded consistently. **A–C,** Residence. Difference vectors link an agent/group to an associated place (e.g., “*doctor*”→“*hospital*”); CCGP is significant. **D–F**, Protectors. Vectors link a protector/guide to the protected individual/group (e.g., “*teacher*”→“*grader*”); CCGP is significant. **G–I**, Antonyms. Antonym vectors are oriented according to valence; CCGP is significant. See Supplementary Figure 3 for a full list of word-pair labels in the MDS plot, color-coded consistently. **J–L**, Verb negation. Vectors connect affirmative verbs to their negated forms (e.g., “*laugh*”→“*not+laugh*”); CCGP is significant. **M–O**, Noun plural. Vectors connect singular nouns to their plural forms (suffix -s); CCGP is significant (p(label) < 0.05; p(neuron) > 0.05). **P–R**, Adjective comparative. Vectors connect positive adjectives to comparative forms (e.g., “*tall*”→“*taller*”); CCGP is significant. **S,** Testing whether the adjective comparative axis reflects a generic -er morphology: mean cosine similarity of derivational agentive -er pairs (e.g., “*teacher*”−“*teach*”, “*keeper*”−“*keep*”) to the adjective comparative pair directions, shown against a random-pair null (grey violin). **T,** Testing whether the noun single/plural axis can be generalized to a verb +s axis: mean cosine similarity of third-person singular verb vs base-form pairs (e.g., “*pulls*”−“*pull*”) to the noun plural–single pair directions. **U,** Predicting deviation from parallel of 206 individual word pairs from the corresponding deviation in GPT-2 embeddings. For visualization purposes, HPC cosine similarities shown here were corrected for 15 individual analogy types as random intercept and random slopes according to the linear mixed effect model, see Results. Showing a scatter plot, linear fit line and correlation coefficient. All significant aligned units (see Supplementary Figure 2 for percentage) in GPT-2 (1280 total units per layer) or BERT (768 total units) were used for this analysis.

We also identified the category of protectors, describing a relationship in which one agent is charged with guarding, protecting, or guiding the other (**Figure 3D-F**). Our examples included “*teacher*”/ “*fifth grader*,” “*doctor*”/“*patient*,” and “*sergeant*”/“*soldier*”. All pairs showed greater alignment than expected by chance, except for “*parents*”/“*child*” (q = 0.34). Overall, the proportion of aligned pairs was much greater than chance (p < 0.0001, binomial test; CCGP: p < 0.05).

We observed the same with all categories. These included **antonyms** (**Figure 3G-I**), such as “*different*”/“*same*”, “*long*”/“*short*,” and “*high*”/“*low*.” (Note that for antonyms, we assigned valence according to Mohammad, 2025). Overall, 12/22 pairs of antonyms were aligned. Unaligned antonyms did not show any obvious pattern and included “*young*”/“*old*” and “*wrong*”/“*right*” (q > 0.05). Nonetheless, the proportion of aligned pairs, while far from complete, was still much greater than expected by chance (p < 0.001, binomial test).

We also tested **verbal negation**, in which the word “*not*” could appear before verbs like “*realize*,” “*laugh*,” and “*happen*” (**Figure 3J-L**). Overall, 13/19 pairs of verbs and their negated forms showed alignment (p < 0.0001, binomial test). Verbal negation has some conceptual similarity to antonymy, although several philosophers and linguists have argued they serve different linguistic functions (Clark & Chase, 1972; Kennedy, 2007; Montague & Thomason, 1978; Quine, 1960). We therefore asked whether the coding direction for antonyms is aligned with the verb negation axis. It is not. Training on antonyms and testing on verb negation resulted in only 2 of the 19 pairs showing significant alignment. This proportion is non-significant (p = 0.24, binomial test, **Supplementary Figure 5A-B**). We observed similar results in the reverse direction: training on verbal negatives and testing on antonyms yielded only 2 of 22 significantly aligned pairs, a proportion that is likewise non-significant (p = 0.30, binomial test).

We did, however, find alignment for two other grammatical operators, the nominal plural marker *- s* (“*boy*”/“*boys*”, **Figure 3M-O**), and the comparative marker *-er*, which changes positive adjectives to comparatives (“*close*”/“*closer*”, **Figure 3P-R**). We found alignment in two other semantic relationships: odd number + 1 (“*one*”/“*two*”) and qualifiers (“*kind of*” +, “*slightly*” +) as well (**Supplementary Figure 3I–P**).

It is important to investigate if the apparent axis alignment arises from any systematic changes in other vector properties—for example, if comparative forms simply differed from their stem adjectives primarily by vector magnitude/length, or if negated verbs were represented as near-perfect inversions (≈180° rotations) of their affirmative forms. They did not. We addressed this by examining the geometry of the *word-pair vectors themselves* (**Supplementary Figure 5C-D**). Because our analysis normalizes vector length prior to computing difference vectors (**Methods**), any residual alignment cannot be attributed to consistent differences in raw vector norms across word forms. Moreover, direct analysis of pairwise vector angles showed that neither verbal negation (e.g., “*grow*” vs. “not *grow*”) nor comparison (e.g., “*tall*” vs. “*taller*”) behaved like simple inversions, consistent with Zuanazzi et al., (2024); instead, these transformations tended to be expressed as directions that are approximately orthogonal to the root-word vectors, rather than as sign flips or uniform length scaling (**Supplementary Figure 5D**). Together, these controls indicate that the semantic axes we identify reflect a genuine direction-consistent relational transformation, rather than a byproduct of systematic norm changes or antipodal structure in the underlying word vectors.

In all six relationships shown in **Figure 3**, as in the gender relationship (**Figure 2**), limiting our control set to the matching part of speech (POS) did not compromise the measured alignment (p < 0.05 in all cases, binomial test). And, not surprisingly, in all cases, we found the same results with BERT no matter if we limit POS or not (p < 0.0001).

We also tested for potential confounds from morphosyntax. Fortunately, the English language offers convenient controls, because the agentive derivational suffix *-er* is identical to the adjective comparative suffix. We identified two word pairs in which our corpus contained both a verb and its derived form (“*teach*”/“*teacher*” and “*keep*”/“*keeper*”). Neither of these was aligned with the comparative adjective axis shown in **Figure 3S**. While there are only two, the chance that these two of them rank the lowest in cosine similarity together with the adj comparative pairs is 2.78% (i.e., p = 0.0278). We conclude that the positive/comparative axis is selective for that relationship; it is not a general -*er* axis.

Likewise, in English, the third person present tense, simple aspect conjugation *-s* (“he *pull**s*** the rope”) has the same morphological marking as the plural *-s* (“the *dog**s*** in the park”). We asked whether verb pairs in the infinitive vs. present simple conjugations are aligned with the noun plural axis shown in **Figure 3M**. They are not (**Figure 3T**). Indeed, only three of the 19 pairs were significant. This proportion is much lower than the proportion of noun plural pairs (23/43) that are significant (p = 0.0051, left-tail Fisher’s exact test).

We have thus far demonstrated that the semantic axes are appropriately parallel, with cosine similarities consistently and significantly positive yet distinct from identity (0 < cos < 1). However, it remains unanswered: what predicts the degree of deviation of individual word pairs from perfect parallelism? We hypothesized that these deviations in the hippocampal (HPC) population are not randomly structured but can be predicted by the deviation of individual words pairs in LLMs. To test this, we constructed a linear mixed effects (LME) model, which was applied to the cosine similarities of individual word pairs across all fifteen diverse analogies. The model was specified as HPC ∼ 1 + LLM + (LLM | analogy type), where we assessed the fixed effect of the LLM similarity while accounting for random intercepts and slopes across analogy types. Our analysis of the 13 layers of BERT failed to reveal any significant alignment with HPC after false-discovery rate correction for multiple layers tested (**Supplementary Figure 5E**). In contrast, analysis of GPT-2’s 37 layers showed robust alignment in middle to late layers (**Supplementary Figure 5F**). For example, HPC cosine similarity was predictable from GPT-2’s last layer (p = 0.025, **Figure 3U**), indicating that hippocampal neural populations might structure the deviations from geometric parallelism in the same way as in some LLMs.

### Semantic axes in pronouns formed prismatic structures

Our corpus contained all seven of the English personal pronouns in both nominative (e.g., “*I*”) and accusative (e.g., “*me*”) forms. Of these seven, we found alignment for five (only the pair “*it (nominative)*”/“*it (accusative)*” and “*you (nominative)*”/“*you (accusative)*” were not aligned (**Figure 4A-C**). This proportion is significant (p < 0.0001, binomial test). Our corpus also contained all seven of the English personal pronouns in the possessive (“*my*”) form. The nominative/possessive forms (“*I*”/“*my*”) were also aligned (p < 0.0001, **Figure 4E-F**). Of the seven accusative/possessive pairs, we found alignment for six (only the pair “*her*”/“*her*” was not aligned, **Figure 4G-I**). This proportion is much greater than chance (p < 0.001). English pronouns also have a singular/plural semantic axis (e.g., “*I*”/“*we*”), which can also appear in the accusative case (“*me*”/“*us*”) and in the nominative possessive form (“*my*”/ “*our*”). All seven examples of the singular/plural set were aligned, p < 0.0001, **Figure 4J-L**.

**Figure 4.**
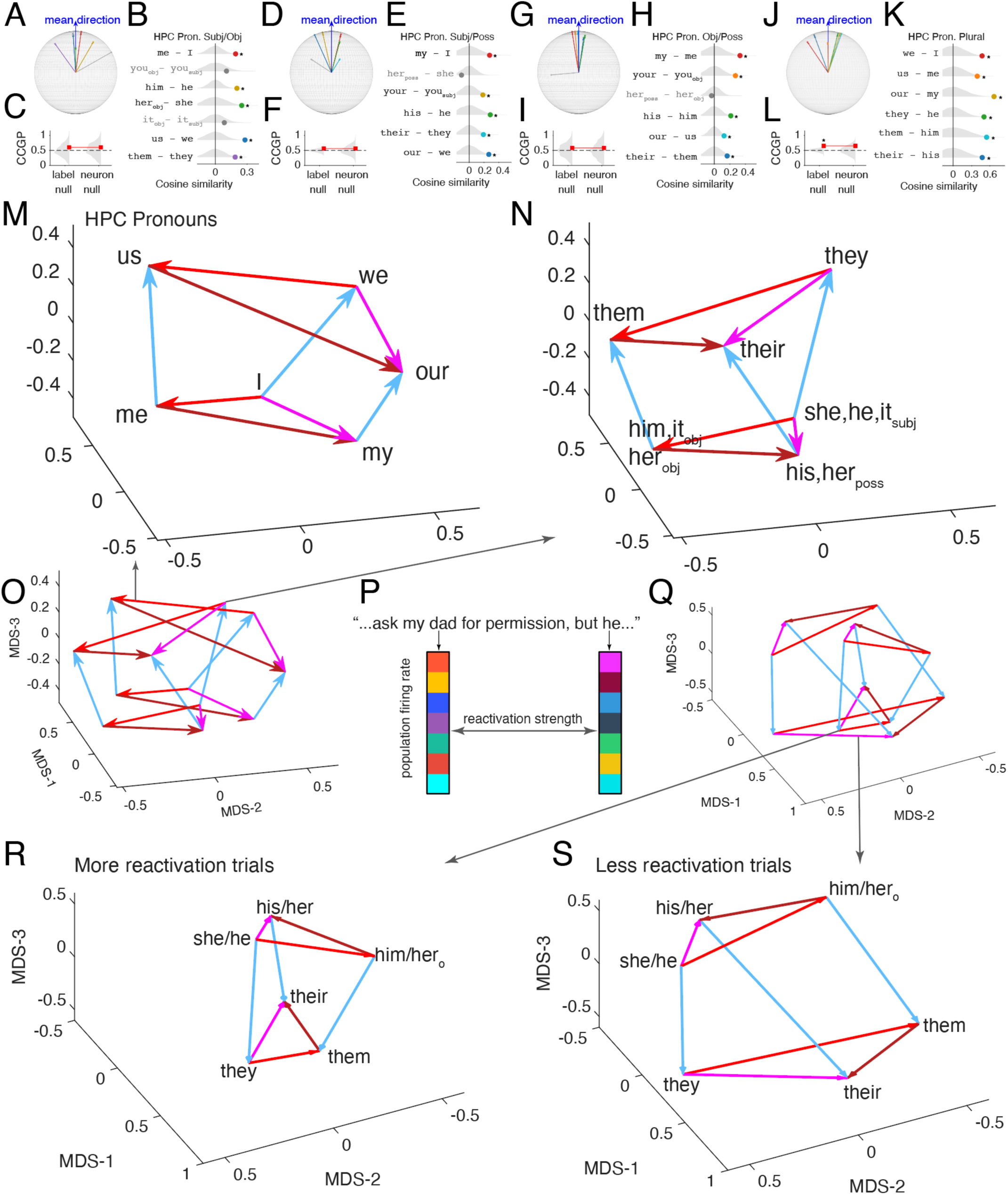
| Hippocampal reactivations consolidate the compositional semantic axes of pronouns. **A–C,** Subject–object case (Pron. Subj/Obj). Difference vectors between nominative and accusative pronoun forms in the hippocampus (e.g., “*I*”→“*me*”) are tail-aligned and visualized on a unit sphere (“globe”; **A**) after a rigid rotation so the within-relation mean direction points upward (blue; visualization only). **B**, For each pronoun pair, the dot shows its mean cosine similarity to the direction of other word pairs computed in the high-dimensional neural population space (not the visualization space); grey violins show the null distribution from random word-pair differences (n = 10000 draws; see **Methods**). Asterisks indicate significant alignment after Benjamini–Hochberg FDR correction across pairs within the relation (q < 0.05). Pairs shown in grey do not reach significance (including “*you*” and “*it*” in nominative vs accusative), yielding 5/7 aligned pairs overall (binomial p < 0.0001). **C,** Cross-condition generalization performance (CCGP) for decoding nominative vs accusative case using a linear classifier trained on a subset of pairs and tested on a held-out pair (see **Methods**). Red markers show observed CCGP; grey violins show label-shuffle and neuron-shuffle nulls (n = 600 shuffles); dashed line indicates chance. * = P < 0.05 **D–F,** Subject–possessive (Pron. Subj/Poss). Same conventions as **A–C**, but for nominative vs possessive determiners (e.g., “*I*”→“*my*”). **G–I,** Object–possessive (Pron. Obj/Poss). Same conventions as **A–C**, but for accusative vs possessive forms (e.g., “*me*”→“*my*”). The homographic possessive/object form “*her*” (poss vs obj) does not show reliable alignment (grey). **J–L,** Number (Pron. Plural). Same conventions as **A–C**, but for singular→plural transformations expressed across case/possessive contexts (e.g., “*I*”→“*we*”, “*me*”→“*us*”, “*my*”→“*our*”, and third-person analogs). All tested singular–plural pairs are aligned (binomial p < 0.0001). CCGP exceeds the label-shuffle null (asterisk) (p(label) < 0.05), but not the neuron-shuffle null. **Related visualizations.** Multidimensional-scaling (MDS) plots of the individual pronoun analogies geometry matching panels **A-L** are shown in **Supplementary** Fig. 6A**-E** **M-O,** Joint pronoun geometry in hippocampal population space. Multidimensional scaling (MDS) visualization of the mean neural population vectors for pronouns spanning the person × number × case triad. Arrows highlight the four component transformations tested in panels **A–L**, using a consistent color code: number (singular→plural; blue), subject→object (nominative→accusative; red), subject→possessive (magenta), and object→possessive (dark red/brown). The lower-left subplot **O** shows the full configuration with all pronoun vertices/edges; the top **M** (first-person) and right **N** (third-person) subplots are the same embedding with the other pronouns removed for clarity (coordinates unchanged). Third-person singular forms are pooled where indicated (e.g., “*she, he, it_subj_”, “him, it*_obj_”, “*his, her*_poss_”). Second person “you” were used for the MDS dimension reduction but not shown due to lack of number (single/plural) designation. Visually, the third-person prism appears more distorted than the first-person prism. n = 150 hippocampus neurons (top 50 most tuned neurons from each of the 4 analogies in panel A-L pooled). **P,** Reactivation quantification. Pearson correlation of population firing rate between each human referring third person pronoun and its antecedents were used to denote the reactivation strength. Same neurons used as in **M-O**. **Q-S,** Same as in **M-O**, but only for third person pronouns, splitting trials into more (**R)** or less (**S)** reactivation. Visually, the third-person prism formed by trials with less antecedent reactivation appeared more distorted, resembled less like a prism than its counterpart with more reactivation.

So far, we tested pronoun analogies one relation at a time. These pairwise tests do not determine whether the full set of pronouns forms a jointly consistent geometry across multiple features, nor do they constrain the relative magnitudes of the transformations. To address this, we analyzed the twelve pronouns spanning the person × number × case, which included first person {“*I*”, “*me*”, “*my*”, “*we*”, “*us*”, “*our*”} and third person {“*he*”, “*him*”, “*his*”, “*they*”, “*them*”, “*their*”}. (We did not have all necessary (single/plural) pronoun type labels for the second person “*you*” and “*your*”, so they were only used for visualization, **Figure 4O, 4P** but not any formal analysis). For statistical power, we pooled third person singular pronouns for this analysis. For each word, we averaged firing rates across trials to obtain a single population vector per pronoun, then z-scored each neuron’s responses across the twelve words. The case × number combinations for the first-person pronouns reveal a clear prism-shaped structure in 3-d space after dimension reduction (**Figure 4M**). Likewise, the case × number combinations for the third person (**Figure 4N**) reveals another prism. Moreover, these prisms largely overlapped but remained visually separable (**Figure 4O**).

To quantify whether these prisms reflect a genuinely compositional structure (Phillips, 2022)—rather than a set of independent pairwise parallels—we used a loop-closure (commutativity) test defined on each 2×2 face of the prism (Andreas, 2019). Consider any quadrangle with four vertices a, b, c, d arranged as `(plural, case1) = a`, `(singular, case1) = b`, `(plural, case2) = c`, and `(singular, case2) = d`. If number and case compose independently (i.e., the representation is locally factorized), then changing number should not depend on case, and equivalently changing case should not depend on number. This can be expressed as the commutator/closure vector `|(a − b) − (c − d)|`, which should be close to zero in an ideal prism because `(a − b)`and`(c − d)` are the same “number shift” computed in two different cases. By simple rearrangement, this is equivalent to `|(a − c) − (b − d)|`, which instead compares the “case shift” at plural vs. singular. Thus, the same statistic tests path-independence: going from singular→plural and then case1→case2 should land in the same place as going case1→case2 and then singular→plural. From the non-reduced, high- dimensional neural vector data we computed this closure error for the three unique case-pair rectangles within each person prism separately (SUB–OBJ, SUB–POSS, OBJ–POSS) and summarized prism distortion (non-closure) as the mean Euclidean norm of these closure vectors; this statistic penalizes both directional and magnitude inconsistencies across the rectangle edges, and goes beyond asking whether any single displacement (e.g., “*we*”–“*I*”) is nonzero or parallel to another displacement. Significance was assessed against a label-shuffled null distribution (500 shuffles).

Overall, loop-closure error across both person prisms was significantly lower than expected under the null (p = 0.002, permutation test), indicating that the joint structure across person/number/case is not explained by chance alignment of word means. When stratified by person, loop closure was robust for first-person pronouns (p = 0.004) but substantially weaker for third-person pronouns (p = 0.084 when considered in isolation). With a matched vertex label swapping permutation test between the first and the third person prism, we further confirmed that the first-person prism was significantly more closed than the third person one (p = 0.0312). This dissociation can be observed in the plots (**Figure 4M-O**): the third-person configuration (**Figure 4N**) appeared more distorted, consistent with reduced commutativity (i.e., the rectangle edges fail to match as closely across case/number for third person). Importantly, this distortion was largely driven by trials with poor antecedent retrieval. We quantified reactivation strength using the Pearson correlation of the population firing rate between each third-person pronoun and its antecedent (**Figure 4P**). When splitting the third-person trials by this metric, loop closure was robust for the half of trials with more reactivation (p = 0.012, **Figure 4 Q-R**), but absent for the half with less reactivation (p = 0.28, **Figure 4S**). A direct comparison with the matched vertex label swapping permutation test confirmed that the prism formed by trials with more reactivation was significantly better closed than its counterpart with less reactivation (p = 0.0156). Overall, these results extend the earlier pairwise parallelogram analyses by showing that at least part of the pronoun system satisfies global closure constraints expected from a prism-like (Trager et al., 2024) compositional (Mitchell & Lapata, 2008, 2010) geometry, rather than only exhibiting local parallelograms.

We next asked whether the same conclusion is supported from a model-based perspective that explicitly tests factorization across features (Kobak et al., 2016). We fit multivariate linear encoding models to the pronoun-elicited neural population responses at individual trial level using (i) a main-effects model with additive terms for person, number, and case and (ii) a full model that also included interaction terms (**Methods**). In the neural data, the full model explained a small fraction of total variance (`R2_full = 0.0175`), indicating substantial neural variability in naturalistic listening that could plausibly reflect (among other factors) context-dependent processing and predictive coding highlighted in prior work (Goldstein et al., 2022; Jain & Huth, 2018; Kumar et al., 2024; Heilbron et al., 2022). Importantly, however, the main-effects component accounted for a large fraction of the variance that was explainable by the full model: the ratio `R2_main/R2_full` was 50.4%, and this fraction was significantly larger than in the label-shuffled null (p = 0.006). Thus, even though the total predictable variance is modest, the predictable component is disproportionately organized along separable axes corresponding to person/number/case. This pattern suggests that the pattern differences associated with each feature are relatively invariant across the other features (i.e., weak interactions), consistent with an approximately separable / partially factorized component in pronoun (Soto et al., 2018; Pooresmaeili et al., 2010) representation. Together, the loop-closure and additive-model results indicate that the pronoun code is not merely a set of independent analogies; it includes a structured component that supports joint composition across features, albeit with person-dependent departures from ideal factorization that are most evident in third person.

Finally, we repeated the same analyses on BERT embeddings for the same pronoun set in the same story contexts (**Supplementary Figure 6F**). In BERT, the low-dimensional embedding exhibited the same prism-like organization found in the neural data, but with more clearly separated first- and third-person prisms. Quantitatively, loop closure was observed, just as it was with neural data (p = 0.002). In the additive-model analysis, BERT showed substantially higher overall explainable structure (`R2_full` = 0.2874), and a stronger degree of factorization: `R2_main/R2_full` was 78.6%, also significant relative to the shuffled null (p = 0.002). Thus, BERT implements a strongly separable, idealized encoding of person/number/case.

### Analogy type dependent semantic axes specialization in ACC and OFC

We next analyzed data from two other brain regions, ACC and OFC. We used neuron-count-matched random subsampling (500 iterations) to account for smaller numbers of neurons in these regions (**Figure 5A**). We found that alignment scores were largely similar to hippocampus in ACC, but a bit lower in OFC (**Figure 5B and C**). In ACC, we found that 13 of the 15 analogies showed alignment, while 10 of the 15 analogies showed alignment in OFC.

**Figure 5.**
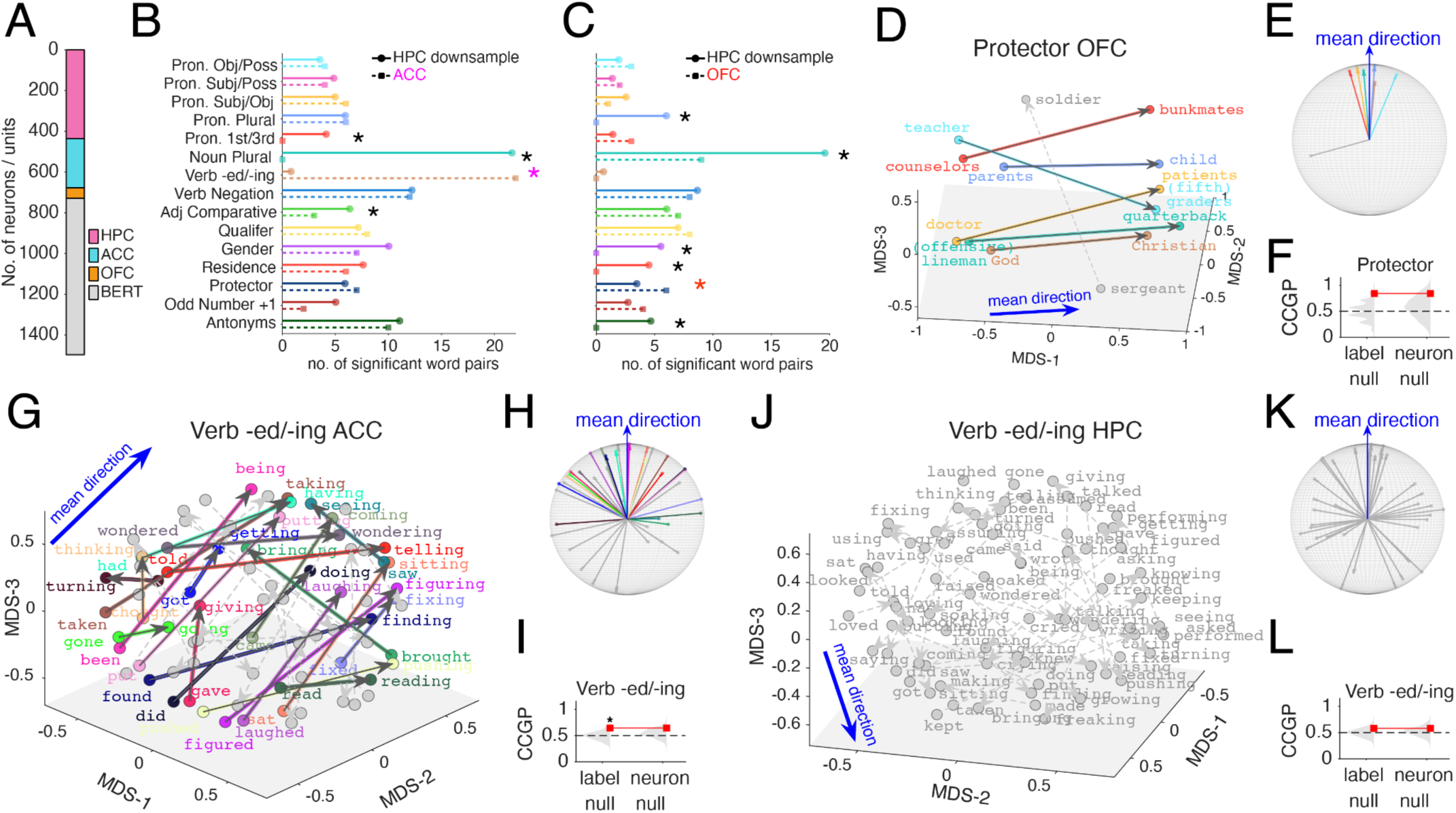
| Complementary regional specialization of hippocampus against anterior cingulate cortex and orbitofrontal cortex across semantic relation types. A, Number of units included from hippocampus (HPC), anterior cingulate cortex (ACC), and orbitofrontal cortex (OFC) along with the number of BERT units/dimensions used for comparison. B, Number of significant aligned word pairs for each analogy type in downsampled HPC (solid circles) and ACC (dashed squares). Symbols denote analogy types with significant HPC–ACC differences after Benjamini–Hochberg FDR correction (Wald F-tests, Satterthwaite degree of freedom method ; q < 0.05; see main text). Significant asterisk color indicated which region had higher value (black, HPC; magenta, ACC) C, Same as in B but for OFC. D–F, Protector relationship in OFC. D, MDS visualization of OFC word embeddings and within-category difference vectors (grey indicates non-significant pairs). Blue arrows denote the within-category mean direction. Grey planes are visual aids only. Colors in D and E match each other. E, Tail-aligned unit-sphere visualization of the same vectors, rotated so the mean direction points upward and normalized in length (visualization only). F, Cross-condition generalization performance (CCGP; red) compared to label-shuffle and neuron-shuffle null distributions (grey violins; dashed line indicates chance). G–I, Verb −ed/−ing relationship in ACC, shown as in D–F. See Supplementary Figure 7 for a full list of word pair labels in MDS plot color-coded consistently. J–L, Verb −ed/−ing relationship in HPC, shown as in D–F.

We assessed areal differences using a brain-area label-shuffling test: we computed the observed cosine similarity under neuron-count matching, then generated a null distribution by pooling neurons across areas, randomly permuting the HPC/ACC label assigned to each neuron (500 shuffles), and repeating the same neuron-count-matched cosine-similarity computation for each shuffle. This non-parametric test yielded p = 0.016, meaning that under random reassignment of neurons to HPC/ACC labels, only 1.6% of shuffles produced an HPC–ACC analogy-type distribution that was at least as divergent as the one observed. Thus, even when controlling for neuron count, the distribution of significant word-pair counts across analogy categories shows an area-specific structure, consistent with specialization of analogy-type selectivity between HPC and ACC. This across analogy area specialization was also observed between HPC and OFC, p = 0.002.

To pinpoint *where* this area-specific structure arises at the word-pair level, we summarized each word pair with a 0–1 significance-prevalence score (sigPrev) that captures how likely that pair is to be deemed significantly aligned with other pairs in HPC in an analogy under ACC-matched sampling (i.e., in HPC, a higher value indicates a more reliably detectable significant word-pair under neuron count matched random subsampling; In ACC, it means the word pair is significant). We fit a linear mixed-effects model with fixed effects of (brain) Area, (analogy) Type, and their interaction, and a random effect for word-pair identity: sigPrev ∼ 1+Area×Type+(1∣PairID), n = 412 word pair observations. Marginal ANOVA tests indicated significant effects of Area *F*(1, 382) = 7.22, p = 0.008, Type *F*(14, 382) = 15.53, p = 7.6e-30 and a robust Area × Type interaction *F*(14, 382) = 9.67, p = 1.8e-18. Thus, regional differences in the reliability/prevalence of significant word-pairs depended on analogy type (highly significant interaction terms (Wischnewski & Peelen, 2021)). The Area × Type interaction was also significant when repeating the analysis with HPC and OFC *F*(14, 382) = 5.68, p = 4.8e-10.

Given the significant interaction, we ran follow-up simple-effects contrasts (Wald F-tests) comparing HPC vs ACC within each analogy type (15 types) and controlled for multiple comparisons using Benjamini–Hochberg FDR correction. After FDR correction, significant HPC–ACC differences were observed for 4 of 15 types: Verb *-ed*/*-ing*: q = 5.5e-11 (more in ACC), Pronoun 1st/3rd: q = 0.0016 (more prevalent in HPC), Noun Plural: q = 9.0e-11 (more in HPC), Adj Comparative: q = 0.024 (more in HPC). All other analogy types showed no reliable HPC–ACC differences after FDR correction (q > 0.05). Similarly, significant HPC–OFC differences were observed for 6 of 15 types: Protector: q = 0.046 (more prevalent in OFC), q < 0.05 more prevalent in HPC for Pronoun Plural, Noun Plural, Gender, Residence, Antonyms. All other analogy types showed no reliable HPC–OFC differences (q > 0.05).

**###Figure 5D-F** shows examples of an analogy type in which OFC shows more prevalent relational structure than the hippocampus (HPC), and a final case in which ACC clearly outperforms HPC (**Figure 5G–L**). For the Protector relationship (e.g., “*sergeant*”/“*soldier*”, “*teacher*”/“*(fifth) graders*”, “*doctor*”/“*patients*”), OFC showed a coherent direction with multiple aligned pairs (colored vectors in **Figure 5D**; concentrated directions in the tail-aligned visualization in **Figure 5E**). This relational geometry is yet to support robust generalization: CCGP was not well above chance, potentially due to the low neuron count in OFC (**Figure 5F**).

Finally, we observed a striking dissociation for the verb *−ed*/*−ing* relationship (past vs. progressive aspectual forms; e.g., “*laughed*” vs “*laughing*”, “*did*” vs “*doing*”, “*told*” vs “*telling*”). In ACC, these vectors exhibited strong within-category alignment (**Figure 5G–H**) and supported above-chance generalization against label shuffle null (**Figure 5I**). In contrast, the hippocampal population did not show a comparably consistent axis for the same verb −ed/−ing pairs (**Figure 5J–K**), and CCGP remained near chance with no reliable improvement over either null distribution (**Figure 5L**). This side-by-side comparison (**Figure 5G–L**) provides an intuitive visualization of the region-level specialization: ACC expresses a much clearer past/progressive axis than HPC, whereas HPC shows advantages for other semantic categories such as pronoun person and noun nominal number.

### Partial neuron-level functional specialization for analogies

We next asked whether the coherent semantic axes we found are mediated by specialized *analogy neurons* or reflect mixed-selective semantic coding (Rigotti et al., 2013; Fusi et al., 2016). We therefore quantified how single-neuron tuning generalizes across analogy types and how population readouts re-weight neurons across analogy types (**Figure 6**). Throughout, we use the term *across-class* tuning to refer to the original class decoding for a given analogy (e.g., gender, **Figure 1E**), summarized by per-neuron area-under-the-curve (AUC, computed from firing-rate based decoding within each analogy, **Figure 1E** and **Methods**). Analyses were repeated under two inclusion regimes: (i) significantly selective neurons only (p(AUC) < 0.05) and (ii) all neurons. The same analyses were performed on hippocampal neurons and on units in the BERT embeddings (paired panels: hippocampus in the upper rows and BERT directly below; e.g., **Figure 6B–D** vs **Figure 6E–G**), as well as in ACC and OFC (**Supplementary Figure 8**).

**Figure 6.**
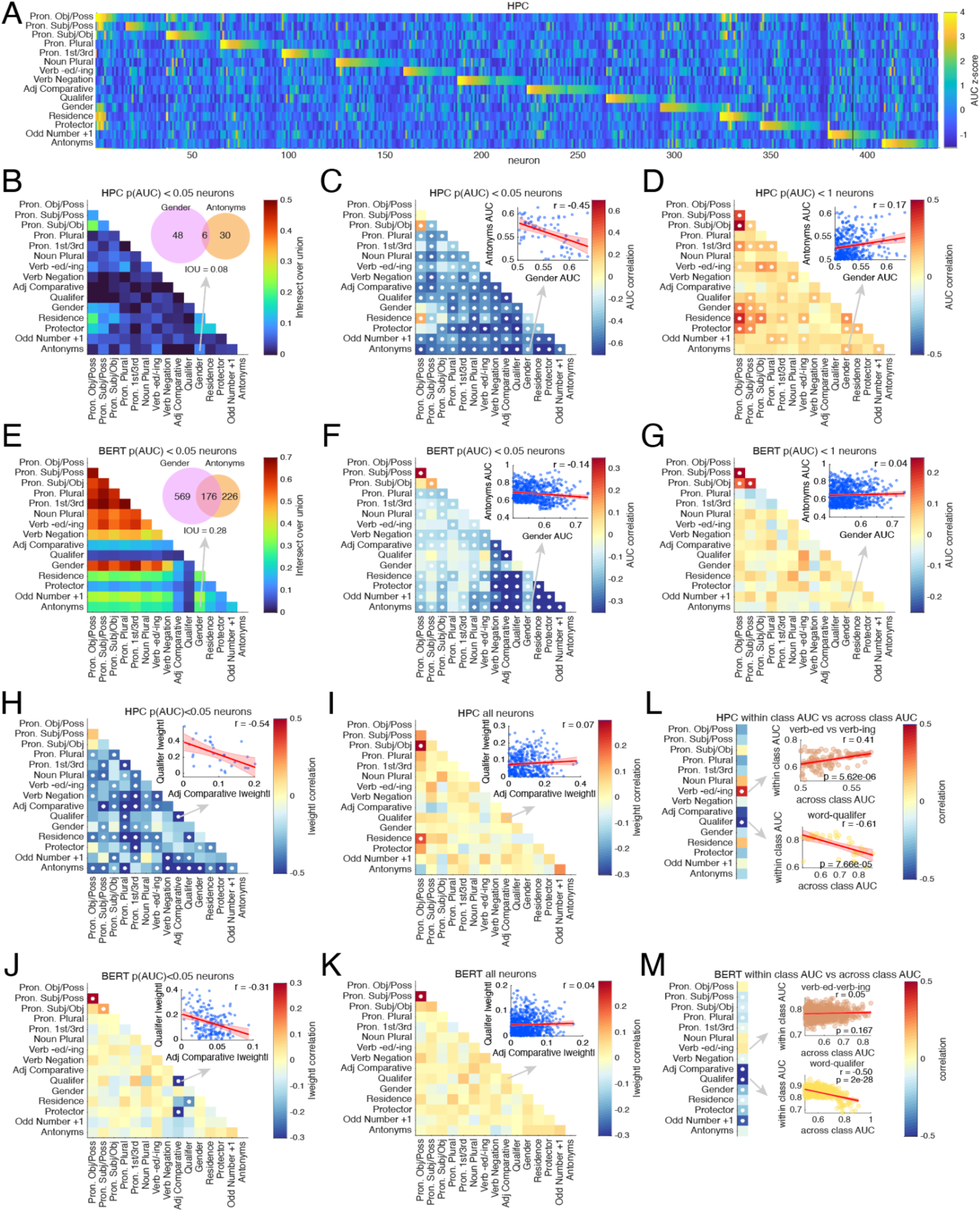
| Functional specialization of semantic axis coding neurons. **A,** Across-class semantic direction coding in hippocampus (HPC). Heat map shows per-neuron decoding strength (AUC values z-scored per analogy type) for each of 15 analogy types (rows) across all recorded hippocampal neurons (columns; n = 437). Neurons are sorted by their most strongly tuned analogy type, producing a near-diagonal block structure consistent with partial factorization. **B,** Overlap of tuned-neuron sets across analogy types in HPC. Each cell shows the intersection-over-union between the sets of neurons significantly tuned to each pair of analogies significance assessed via the label-shuffled null described in Fig. 1E and **Methods**). Inset: example overlap for Gender vs Antonyms (48 and 30 tuned neurons, 6 shared; IoU = 0.08). **C,** Pairwise coupling of tuning magnitudes across analogy types in HPC among tuned neurons. For each analogy pair, the color indicates the Pearson correlation (r) between neurons’ AUC values computed for the two analogies, restricted to the union set of tuned neurons for either analogy (p(AUC) < 0.05 for at least one member of the pair). White dots indicate correlations significant after Benjamini–Hochberg FDR correction across analogy pairs (q < 0.05). Inset: Gender AUC vs Antonyms AUC, each dot is a neuron, red line and shade is linear regression fit and 95% CI. **D,** Same as **C**, but including all hippocampal neurons. Correlations shift toward weakly positive values, indicating weak shared structure when the full population is included. **E–G,** Same analyses as **B–D**, performed on BERT embedding “units” (last-layer dimensions; 768 units;). **E,** IoU overlap matrix for tuned BERT units. inset: Gender vs Antonyms (569 and 226 tuned units, 176 shared; IoU = 0.28). **F,** AUC–AUC correlation matrix among tuned units (white dots: FDR-significant; inset r = −0.14). **G,** AUC–AUC correlations across all units (inset r = 0.04). **H–K,** Factorization at the level of optimal linear population readouts. For each analogy type, a logistic-regression decoder was fit, and each neuron/unit’s contribution was summarized by the absolute decoder weight |w|. Heat maps show pairwise Pearson correlations between |w| vectors across analogy types. **H,** HPC tuned neurons, with predominantly negative correlations; white dots indicate FDR -significant correlations (q < 0.05). Inset: Adj Comparative vs Qualifier |w| (r = −0.54). **I,** HPC all neurons, with correlations near zero/weakly positive (inset r = 0.07). **J,** BERT tuned units (inset r = −0.31). **K,** BERT all units (inset r = 0.04). **L–M,** Relationship between across-class and within-class coding. For each analogy type, within-class AUC (mean discriminability among words within the same side of the analogy) was compared to the original across-class AUC across neurons/units; the left column summarizes the correlation for each analogy type (color indicates the correlation coefficient). Insets show example relationships. **L,** HPC: within-class vs across-class AUC correlations can be positive (verb-ed vs verb-ing: r = 0.41, p < 0.0001) or negative (word–qualifier: r = −0.61, p < 0.0001), indicating partial separation of axis coding from within-side “role/identity” coding. **M,** BERT: weak coupling for verb-ed vs verb-ing (r = 0.05, p = 0.167) but strong negative coupling for word–qualifier (r = −0.50, p < 0.0001).

After sorting hippocampal neurons by their most strongly tuned analogy type, the AUC matrix exhibited a near-diagonal block structure, suggestive of specialization (**Figure 6A**). Specifically, each analogy type was associated with a distinct set of neurons that had elevated participation in that analogy type, indicating that a partially discrete subsets of neurons preferentially support different analogy dimensions. Importantly, however, there were also moderate off-diagonal activations, indicating that many neurons had above-baseline tuning for analogy types beyond their preferred one. Thus, semantic codes are partly multiplexed.

To quantify these observations, we used an intersection-over-union approach (IoU; Jaccard index, Jaccard, 1912) between the sets of neurons significantly participating in each analogy pair (**Figure 6B**). In the hippocampus, IoUs were generally low across most analogy pairs, indicating that significant tuning is typically carried by distinct subpopulations for different analogies. The inset example (gender vs. antonyms) illustrates this low overlap (6 shared neurons out of 48 and 30 tuned neurons, respectively; IoU = 0.08). Interestingly, analogy pairs involving closely related, human-referential analogies (e.g., pronoun-related and social-role analogies) tended to show comparatively higher IoU values than unrelated pairs, suggesting partial sharing within semantically coherent clusters. In contrast, BERT exhibited substantially larger overlaps between tuned unit sets across analogy types (**Figure 6E**). For the same illustrative pair, gender and antonyms included 569 and 226 tuned units with 176 shared (IoU = 0.28), consistent with more widespread reuse of features across analogy types, potentially reflecting the distributed mixing of information across embedding dimensions produced by the model’s repeated self-attention and densely connected feed-forward sublayers throughout the transformer (Clark et al., 2019; Devlin et al., 2019).

Overlap alone does not determine whether neurons that participate in multiple analogies do so with similar *strength*. We therefore asked, among neurons tuned to at least one of two analogy types (union set: p(AUC) < 0.05 for either member of the pair), whether a neuron’s AUC for one analogy predicts its AUC for another. In the hippocampus, pairwise correlations of AUC across analogy types were predominantly negative (**Figure 6C**; significant analogy type pairs marked after FDR correction), indicating specialization. In other words, neurons that strongly separate the two classes within one analogy tend to have weaker roles for other analogy types. The inset in Figure 6C (gender vs antonyms) shows an example negative relationship (r = −0.45). A limited set of positive correlations appeared, most notably among closely related pronoun-based analogies and select human-referential categories. BERT showed the same tendency under the tuned-unit restriction but with smaller magnitude correlations (**Figure 6F**), indicating that even when many units are significant for multiple analogies (high IoU, **Figure 6E**), tuning magnitude can still display specialization.

When we expanded the analysis to include all neurons, the AUC–AUC correlation structure shifted: correlations became weakly positive across much of the matrix in HPC (**Figure 6D**) and, similarly, moved toward near-zero/slightly positive in BERT (**Figure 6G**). This reversal suggests a two-tiered organization: specialized strongly selective neurons overlaid on a less specialized set of less selective generalists.

To test whether the same principles hold for population coding, we fit a logistic regression model decoding member classes (e.g., male vs female in the gender analogy) of each analogy type from neural population activities and decoder used the absolute weight magnitude (|w|) assigned to each neuron(brain)/unit(BERT) as an index of its contribution to that analogy’s population decision variable. We then computed correlations between |w| vectors for each analogy pair. In the hippocampus, when restricting to tuned neurons, weight correlations were negative (**Figure 6H**), paralleling the AUC anti-correlations and showing that factorization persists at the level of the optimal linear readout (t-test for a correlation coefficient, q < 0.05 after FDR correction). The inset illustrates an example of strong negative correlation of weight vectors (Adj Comparative vs Qualifier; r = −0.54), indicating that decoders (or potential downstream neural readouts) for different analogy types tend to recruit the subsets of the tuned population at starkly divergent strengths. In BERT, tuned-unit weight correlations were generally closer to zero, with fewer significant effects (**Figure 6J**; example r = −0.31), again consistent with a less sharply segregated readout structure. When all neurons/units were included (hippocampus: **Figure 6I**; BERT: **Figure 6K**), weight correlations moved toward near-zero or weakly positive values (HPC example r = 0.07; BERT example r = 0.04), consistent with broadly distributed and orthogonalized contribution plus a tiny, shared component rather than sharply task-specific recruitment across the neural population.

Finally, we asked whether neurons that encode an analogy’s (across-class separation) also encode *within-class* distinctions among words on the same side of the analogy—operationally defined as “role” coding within-class (e.g., “*man*” vs “*king*” within the male set; “*woman*” vs “*queen*” within the female set). They mostly do not. For each analogy type, we computed a within-class AUC (averaged over within-class pairwise discriminations) and correlated it across neurons with the original across-class AUC for that analogy (**Figure 6L–M**). Within high tuning strength (low p(AUC)) neurons, within-class and across-class tuning were often not or weakly related and could be either positively or negatively coupled depending on analogy type (**Figure 6L**). For verb tense (verb-*ed* vs verb-*ing*), within-class AUC was positively correlated with across-class AUC (r = 0.41, p < 0.0001), whereas for word–qualifier the relationship was strongly negative (r = −0.61, p < 0.0001). The prevalence of negative (and sometimes significant) correlations indicates that across-class information and within-class role/identity information are at least partly factorized across neurons: neurons best at separating the two sides of an analogy are not necessarily those best at discriminating items within a side. BERT also showed a strong tendency toward factorization for certain analogy types (**Figure 6M**), including a robust negative relationship for word–qualifier (r = −0.50, p < 0.0001), while verb tense showed little coupling (r = 0.05, p = 0.167). We observed qualitatively similar specialization effects in ACC and OFC (**Supplementary Figure 8**).

## DISCUSSION

Our results suggest that semantic information is organized along axes within a high-dimensional neural representational space. These semantic directions are consistent across sets of words that stand in similar relationships, yielding parallelogram-like structures in the embedding space. This geometric structure could provide a crucial mechanistic solution to analogical reasoning, an integral part of human cognition. Analogical reasoning enables us to perform abstraction, knowledge transfer, and learning across superficially different domains through the preservation of higher-order structure (Gentner, 1983; Gentner & Markman, 1997; Hofstadter, 2001; Holyoak & Thagard, 1996; Hofstadter, 1995; Gentner, 2010). By representing semantic relationships as consistent directional vectors, the brain can map relational structures and solve analogies through processes akin to vectorial arithmetic (Gärdenfors, 2000, 2014; Mikolov et al., 2013; Rumelhart & Abrahamson, 1973). Furthermore, this vector-based geometry provides a potential neural substrate for grammatical productivity, consistent with speakers’ ability to apply morphological rules to novel items (e.g., Berko, 1958).

While previous neuroimaging studies have provided macroscopic evidence for semantic directions in the brain, investigating the true geometric structure of these representations has been bottlenecked by the spatial and temporal resolution of fMRI. For instance, hemodynamic responses in fronto-temporo-parietal networks exhibit vector-arithmetic properties during highly controlled tasks (Wu et al., 2022), and brain-wide encoding models have identified coarse semantic axes during naturalistic listening (Zhang et al., 2020). However, critical gaps remain. Analyses relying on encoding models and pooled subject data make it difficult to disentangle genuine neural relational geometry from the inductive biases and assumptions of the models themselves (Popov et al., 2018; Kriegeskorte & Douglas, 2019). More critically, because fMRI voxels aggregate responses of thousands of neurons in a 1–10 mm^3^ volume over seconds, disentangling the semantic specialization of individual neurons during rapid natural speech (∼250 ms per word) proves highly elusive. Our results bypass these limitations to provide a precise, neural population-level view of semantic coding. We go beyond demonstrating the mere existence of semantic axes. By examining the fine-scale relational geometry of these neural populations, we reveal that deviations from perfect parallelism are not unstructured or random noise. Rather, these nuanced geometric deviations can be directly predicted by the semantic representations learned by LLMs, establishing a powerful structural link between biological and computational semantic spaces.

The near-closed prism geometry for pronouns—spanning person × number × case —supports the view that these grammatical features contribute an approximately separable (invariant) component to the pronoun code—i.e., the displacement associated with one feature is largely preserved across the levels of the others (Ashby & Townsend, 1986; Soto et al., 2018), consistent with a transformation-based view of factorized representations (Higgins et al., 2018). The prism’s near-closure further implies approximate path-independence, such that applying number- and case-related transformations in either order yields similar endpoints, consistent with commuting transformations in symmetry-based accounts of factorized representations (Higgins et al., 2018; Zhu et al., 2021). By contrast, a purely pairwise association / lookup-table scheme—or an exemplar-based account in which each mapping is learned independently—does not, without additional compositional structure, naturally predict systematic closure and order-invariant composition across multiple dimensions (Nosofsky, 1986). Importantly, this kind of closure is not guaranteed by high-dimensional representations in general; rather, these systematic order-invariant composition is a diagnostic signature of structured factorization/disentanglement, which typically requires specific inductive biases or architectural constraints (Higgins et al., 2022; Locatello et al., 2020; Higgins et al., 2018), raising the possibility that the brain operates under comparable compositional constraints. More broadly, the emergence of such factorized structure at the level of neural population geometry suggests a concrete mechanism by which the brain could support rule-like generalization over linguistic variables without invoking discrete symbolic representations (Smolensky, 1988). Such an organization naturally supports productivity, in that a small set of learned operators can be recombined to generate a large space of grammatical forms, and it suggests a learning advantage whereby new combinations can be inferred from existing structure rather than acquired through exhaustive experience (Berko, 1958; Courellis et al., 2024; Flesch et al., 2022, 2023; Kemp & Tenenbaum, 2008).

Meanwhile, during inferential reasoning, single units across the hippocampus simultaneously encode multiple variables in a factorized format, and that this geometry emerges specifically after learning to perform inference (Courellis et al., 2024). Our results confirm these important results, and extend them to uncued, spontaneous representations, to the naturalistic listening context, and to brain regions beyond the hippocampus. Finally in a controlled reading paradigm, pronouns can reactivate hippocampal representations of their antecedent concepts (Dijksterhuis et al., 2024). Our findings extend this view by showing that pronouns occupy a structured feature space in hippocampal population activity (Figure 4).

Our results therefore extend this work by showing that pronoun feature geometry can constrain and delimit candidate antecedents (e.g., by narrowing person/number/case-consistent referents). Crucially, our finding that the third-person pronoun prism successfully closes only during trials with high antecedent reactivation suggests that this structured geometric representation is not unconditionally instantiated but is instead dynamically dependent on the subject’s ongoing cognitive state. It appears that maintaining a prismatic compositional geometry requires the active grounding of the pronoun to its specific referent in working memory—for instance, successfully linking both the nominative "he" and the genitive "his" back to the same underlying entity, such as "father." The distortion during less reactivation might potentially be explained by cognitive interference; for instance, the neural population might remain occupied by the residual processing of intervening lexical items (e.g., representations of other nouns like "flower" or "dog"). Furthermore, general attentional lapses might also prevent the successful antecedent retrieval necessary to maintain the factorized representation. Consequently, this indicates that the brain’s ability to support composition geometry in pronouns may rely on active reference resolution.

Our parallel results in three brain regions suggest a partial division of labor rather than a uniform, cortex-wide “analogy code.” In most analogy types, hippocampus matched or outperformed ACC, it also outperformed OFC similarly, consistent with a large body of work showing that hippocampal circuits support flexible relational codes that extend beyond physical space to non-spatial task variables and abstract manifolds (Aronov et al., 2017; Knudsen & Wallis, 2021; Nieh et al., 2021), including in humans (Courellis et al., 2024). The clearest exception in our dataset was verb morphology: the *-ed*/*-ing* relation was markedly stronger in ACC than in hippocampus. One parsimonious explanation is functional: dorsal ACC sits on the medial wall within a network of cingulate motor areas that project to motor cortex (and, in primates, have direct spinal projections), and ACC neurons are widely implicated in linking actions to outcomes and maintaining control-relevant task states—computations naturally engaged by verb/event representations and sequential structure (Picard & Strick, 1996; Hayden et al., 2010; Heilbronner & Hayden, 2016). Conversely, OFC showed comparatively strong structure for the protector relation after matching neuron counts (**Figure 5C**), plausibly reflecting OFC’s role in representing latent “task states” and incorporating social context into value-related representations (e.g., who benefits, who is harmed, who is responsible), as well as its sensitivity to socially defined signals/categories (Wilson et al., 2014; Azzi et al., 2012; Watson & Platt, 2012).

Surprisingly, we found some evidence for specialization, in that neurons that most strongly participate in one analogy type are relatively less likely to participate in another. Mechanistically, specialization can emerge when the task family itself decomposes into reusable sub-computations that a network can allocate to partially separable internal machinery. In multitask recurrent neural networks, training one network on many cognitive tasks yields *functional clusters* of units that become specialized for different processes (Yang et al., 2019). Driscoll et al. (2024) similarly found that multitask training can produce modular computational strategies, described as reconfigurable dynamical motifs that mirror the subtask structure of the training set. Given that language is inherently multi-constraint and compositional (with systematic reuse of morpho-syntactic operations and relational templates), it is plausible that the same pressures could bias learning—biological or artificial—toward forming specialized subpopulations that are straightforward to recruit and recombine across related relational demands (Behrens et al., 2018; Lake et al., 2017).

BERT possesses a much higher density of aligned units (**Figure 1F**) and substantially larger overlaps between tuned sets across analogy types compared to the hippocampus (**Figure 6E**). This dense, distributed coding likely stems from the standard transformer architecture, which relies on dense mixing across hidden dimensions and all-to-all token interactions during self-attention (Devlin et al., 2019; Vaswani et al., 2017). Strikingly, BERT embeddings exhibit the same functional specialization that the brain does (**Figure 6F**). This is consistent with a large literature showing that linguistic information is staged across layers (surface/syntax earlier; higher-level semantics and discourse later, Jawahar et al., 2019; Tenney et al., 2019; Rogers et al., 2020), and that only a subset of attention heads carry stable, linguistically interpretable roles (e.g., dependency relations and coreference; Clark et al., 2019; Voita et al., 2019). In further support of this view, our layer-wise scans of both BERT and GPT-2 (Radford et al., 2019; **Supplementary Figure 2**) show that the fraction of embedding dimensions meeting our analogy-tuning criterion generally decreases with layer depth, suggesting progressive compression into fewer, more selectively engaged features.

Importantly, explicit specialization in LLM is *instrumentally useful* for scaling and capability for LLM. Shazeer et al. (2017) and Fedus et al. (2022) showed that a mixture-of-experts architecture can achieve >1000× increases in model capacity with only minor losses in computational efficiency, this is because routing each input word through only a *small subset* of specialist subnetworks allows the overall system to accumulate diverse functions without forcing every parameter to participate in every computation. They reported that different experts become highly specialized in ways that track syntax and semantics. In that light, our specialist analogy-aligned subpopulations could be interpreted as “expert-like” neural resources: a way to store and access many relational skills efficiently, potentially favored in brains because it increases representational capacity and behavioral flexibility under tight metabolic and continual-learning constraints. In other words, the LLM results provide a normative explanation for why the brain would show aligned and factorized representations.

## Acknowledgements

We thank Victoria Gates and Raissa Mathura for invaluable assistance.

## Data availability

The data that support the findings of this study are available from the corresponding authors upon reasonable request.

## METHODS

### Human intracranial neurophysiology

Recordings of 11 adult patients (6 males, 5 females) were collected in the Epilepsy Monitoring Unit (EMU) at Baylor St. Luke’s Hospital, following established intracranial recording procedures during epilepsy monitoring (see also Xiao et al., 2024; Franch et al., 2025). Recordings of 3 patients were collected in the University of Utah Hospital (3 females).

Single-neuron activity was obtained using stereotactic depth electrodes (sEEG), specifically AdTech Medical Behnke–Fried–style probes. On average, each participant had approximately three probes ending in the bilateral hippocampi. Twelve patients had approximately two probes in the bilateral ACC. Five patients had approximately one probe in the OFC. Electrode placement was confirmed by coregistering pre-operative MRI with post-operative CT.

Each depth probe carried eight channels optimized for recording unit activity. Signals were acquired with a 512-channel Blackrock Microsystems Neuroport system at 30 kHz. Spike detection and sorting were performed using Wave_clus (Chaure et al., 2018), followed by manual curation. Units (single- or multi-unit) were retained based on waveform stability and morphology (e.g., slope, amplitude, trough-to-peak features) and inter-spike-interval distributions consistent with a refractory period (no ISIs shorter than 1 ms). Unless otherwise noted, analyses included both single- and multi-unit activity (Franch et al., 2025).

### Electrode localization

Electrode localization followed a standardized imaging workflow (Franch et al., 2025). Briefly, contacts were reconstructed and visualized using iELVis (Groppe et al., 2017), with group-level plotting performed via RAVE (Magnotti et al., 2020). For each participant, pre-operative T1-weighted anatomical MRI and post-operative Stealth CT DICOM images were collected and converted to NIfTI format (Li et al., 2016). The post-operative CT was then aligned to the MRI coordinate space using FSL (Jenkinson & Smith, 2001; Jenkinson et al., 2002).

The aligned CT was imported into BioImage Suite (v3.5β1; Joshi et al., 2011), where electrode contacts were manually identified. Contact coordinates were then transformed into each participant’s native space using iELVis MATLAB utilities (Yang et al., 2012) and displayed on cortical surfaces reconstructed with FreeSurfer (v7.4.1; Dale et al., 1999). Microelectrode locations were defined relative to the first (deepest) macro-contact of the Behnke–Fried depth electrode assembly. Finally, RAVE was used to warp participant anatomy and electrode coordinates into MNI152 space to enable across-subject visualization.

### Natural language stimuli

Participants listened to a set of naturalistic spoken narratives selected to be engaging and linguistically diverse (Franch et al., 2025). Specifically, the stimulus set comprised six episodes from The Moth Radio Hour (each approximately 5–13 minutes long), totaling 47 minutes and 25 seconds of audio (7,346 words). In HPC and ACC analysis, 34 words not heard by all patients with recordings in those two areas were removed. The stories were: “Life Flight,” “The One Club,” “The Tiniest Bouquet,” “My Father’s Hands,” “Wild Women and Dancing Queens,” and “Juggling and Jesus.” Each excerpt featured a single speaker delivering an autobiographical story to a live audience. Audio was presented continuously through the built-in speakers of the participant’s hospital television. To synchronize stimulus timing with neural recordings, the audio output from the playback computer was routed as an analog input directly into the Neural Signal Processor (sampled at 30 kHz), ensuring precise temporal alignment between the stimulus waveform and recorded neural activity.

### Audio transcription

After data collection, the stimulus .wav audio file was transcribed automatically using a Python pipeline with AssemblyAI (Franch et al., 2025). The resulting word-level transcript and timestamps were converted into a Praat-compatible TextGrid, which was then loaded into Praat for manual review. The original .wav file was also imported into Praat so that spectrograms and timing boundaries could be inspected directly. Word onset and offset times were manually corrected to ensure accurate temporal alignment.

### Semantic embedding extraction from language models BERT

Extraction of BERT embeddings was implemented using “bert-base-cased” (Devlin et al., 2018) via Hugging Face Transformers (Wolf et al., 2020), following the approach in Katlowitz et al. (2025). Because BERT is an encoder-only, masked-language model that natively incorporates bidirectional context, embeddings were generated with an expanding-context method intended to approximate left-to-right (causal) processing.

Specifically, the running text began as an empty string, and one new word was appended at a time (retaining punctuation for context). After each addition, the sequence was re-tokenized with BERT’s tokenizer and passed through the model. A running context of maximum size of 512 tokens (current word and past words) were kept. Although BERT encodes both left and right context in principle, only the hidden states associated with the newly appended word were extracted on each step, and subword-piece vectors were averaged to yield a word-level embedding.

Hidden states from all 13 layers (embedding layer plus 12 transformer layers; 768 dimensions per layer) were saved, with analyses focusing on the final layer (layer 12). The implementation used Python with PyTorch (Paszke et al., 2019).

### GPT-2

A parallel embedding pipeline was derived from the transcript using the GPT-2 “gpt2-large” model (Radford et al., 2019) accessed through Hugging Face Transformers (Wolf et al., 2020), following the procedure described in Katlowitz et al. (2025). The transcript spreadsheet was organized so that each row represented a single token, including punctuation and end-of-sentence markers (e.g., “.”, “?”, “!”).

Punctuation was kept in the running text to preserve context, while being tracked to prevent token–word alignment issues during later tokenization. The GPT-2 Large architecture contains 37 total layers (an embedding layer plus 36 transformer blocks), producing 1280-dimensional hidden states, and supported contexts up to 1024 tokens.

To respect GPT-2’s autoregressive structure, embeddings were computed incrementally. The input context started empty and then grew one word at a time to the maximum of 1024 tokens. After each word was appended, the current string was tokenized using GPT-2’s byte-pair encoding tokenizer and passed through the model in inference mode (no gradient computation). Hidden states were extracted from all layers; to produce a single vector per transcript word, the vectors corresponding to that word’s sub-word pieces were averaged.

### AUC

We screened neurons for “analogy separability” using only trial-level firing rates and two word-sets representing the two halves of each analogy (A-words and B-words, e.g., male-words vs female-words). Repeated presentations of the same word, and surface variants grouped to the same member (e.g., *man*/*men*), were pooled within their respective halves. Each word had 14.5 ± 1.9 trials (mean + s.e.m. median = 3 trials). The number of trials for a word pair (defined as the minimum of the A or B side word) did not predict whether that pair was significantly aligned with the other pairs in the same analogy. This was determined using a Wald z-test on the logistic regression coefficient for trial count predicting significant alignment. (HPC: p = 0.18; ACC: p = 0.78; OFC: p = 0.26). For each neuron we quantified how well single-trial firing rates distinguished A-words from B-words using an orientation-free balanced AUC (Compute the usual AUC. If it is below 0.5, flip it by subtracting it from 1. Otherwise keep it as-is. The result is always at least 0.5, where 0.5 indicates chance). To assess whether a neuron’s observed balanced AUC exceeded chance given the exact trial counts, we performed a label-shuffle permutation test at the trial level: Shuffle the A/B labels across trials (keeping the same number of A and B trials) R times. Each time, recompute the balanced AUC. Count how many shuffled values are greater than or equal to the observed balanced AUC, add one to that count, then divide by R plus one. That gives a one-sided p-value. We interpret this p-value as a tuning-strength score for analogy separation: smaller values indicate stronger, more reliable trial-level differences between the two halves. Downstream analyses can shortlist neurons by placing a threshold on this scalar, thereby retaining units that contribute consistent signal to subtraction-style population readouts and discarding units whose subtraction would be centered around zero (no firing rate differences) and thus dominated by random noise.

We developed two additional AUC null distributions, p(AUC-within-column) and p(AUC-word-level) and screened a small number of additional tuned neurons; they were not used (e.g., not in Figure 1E-F) unless otherwise specified. The second word-level permutation control, p(AUC-within-column), uses a within-column trial reassignment null: we keep each word’s column label A/B fixed, but within each column separately we randomly reassign individual trials among the words in that column while preserving the number of trials contributed by each word. After this reassignment we recompute each word’s aggregated firing rate (using the same aggregation rule as above) and then recompute the word-level AUC. This null and resulting p(AUC-within-column) asks whether the observed word-level separability depends on the specific mapping between trials and word identities within each column, while controlling for the empirical trial counts per word and the overall pool of trials available within each column.

In the third word-level AUC null p(AUC-word-level), we treat each unique word in the lexicon as a single data point (rather than each trial). For each neuron and each word, we first aggregate across all repeats of that word to obtain one firing-rate estimate per word (the mean of per-trial rates). Each word inherits its class label A/B from the lexicon column it appears in, and we then compute the same orientation-free AUC using these per-word rates. This word-level formulation reduces the influence of uneven repetition counts across words, because a word that appears many times still contributes only one point to the AUC.

### Relational Similarity

Our aim was to test whether the word pairs from two halves of an analogy share a consistent direction in neural population space. For each analogy type we defined two word-sets—“A-words” and “B-words” (for example, male-words vs female-words)—drawn from the podcast vocabulary. Single-trial firing rates were first averaged across repetitions of the same word to obtain one vector per word and neuron. Surface variants that belong to the same member (for example, *man* and *men*) were pooled within the corresponding half by simple averaging. Before forming any differences, each word’s firing rate vector was L2-normalized to unit length so that subsequent computations reflect direction rather than overall response magnitude.

For each row (one A-member paired with its B-member within the same analogy type), we formed a difference vector by subtracting the A-member vector from the B-member vector and then scaling this difference to unit length. To evaluate alignment for a held-out row, we compared its unit difference vector (“test”) to the unit difference vectors from all other rows of the same analogy type (“training”). The observed score for that held-out row was the average cosine similarity between the test vector and each individual training vector.

Statistical evidence was obtained with a per-row permutation test based on random word differences with random words sampled from the entire podcast vocabulary (not just the A/B sets). In each of 10,000 draws we assembled as many random difference vectors as there are training rows in the dataset, normalizing them exactly as done for the training/testing pairs. For a given held-out row, its null distribution was built by taking, from each draw, the same number of random differences as there are training rows and computing the mean cosine similarity between those random differences and the test vector. The p-value equals “one plus the number of null means at least as large as the observed mean” divided by “one plus ten thousand”; this plus-one adjustment avoids zero p-values. While the number of training rows is constant under leave-one-out, each held-out row has a different test vector, so each row naturally receives its own null distribution of similarity (test vector vs random difference vectors) even as we shared the underlying pool of random words. Intuitively, these null mean cosine similarities are expected to center near zero for any test direction. We performed FDR correction on p values for each analogy according to the number of pairs and corresponding number of p values before designating/counting the number of significant word pairs per analogy type.

We searched a list of tuning strength score p(AUC) cutoffs (21 values from 0.01 to 0.35) and selected a cutoff that yielded a high number of significant pairs per analogy. All neurons with analogical tuning stronger than the cutoff are included in the population vector for words in that analogy. This is based on our hypothesis that if a neuron’s firing rate has no difference between A vs B words (AUC close to 0.5, large p(AUC)), it should contribute to zero-centered random noise during the A - B subtraction style analogy which we investigate. A minimum of 8 neurons with the strongest tuning were included when the criterion returned less than 8 neurons. Due to the large number of artificial neurons(units) in BERT compared to any brain regions (Figure 1F, 6E), we did not search by p(AUC) but instead grid searched [15 30 60 120] units in BERT to identify the optimal number of neurons for this analysis. A complementary screening approach (both p(AUC-within-column) and p(AUC-word-level) smaller than 0.01) were used but yielded a nominal number of additional neurons incorporated in the population vector across 15 analogy types (ACC/OFC: 0.07 ± 0.26 neuron mean ± std, max 1; HPC: 0.47 ± 1.1 neuron, max 4).

For Supplementary Figure 5D only: Because neurons can only maintain non-negative firing rates, raw word vectors tend to point into the same region (positive portions) of a space in any dimension/axes and thus can never be more than 90 degrees with respect to each other; To remove this bias, we centered each neuron by subtracting a baseline mean computed from words outside the lexicon before forming the population vector used in Supplementary Figure 5D.

### MDS visualization

MDS was used for visualization only; we constructed a word-by-neuron response matrix by averaging each neuron’s firing rate across all trials for each word after pooling pre-specified surface-form variants (e.g., singular/plural) into a single word identity. We did not impose any A/B grouping for this analysis. To equalize scaling across neurons, we z-scored responses across words separately for each neuron. We then computed pairwise dissimilarities between words using cosine distance on the resulting population vectors and embedded the words into a 3-dimensional space using MATLAB’s mdscale() (metric multidimensional scaling). The embedding was optimized to minimize the standard MDS stress criterion and plotted as a low-dimensional visualization.

### CCGP

We used a cross-condition generalization performance (CCGP) analysis to ask whether neural population activity supports abstract structure that generalizes across matched word pairs, and whether this generalization exceeds what would be expected from either random labels or random geometry of the neural population. This analysis inherited the neuron selection criteria in the relational similarity analysis.

a. Pseudo-population and matched word pairs. For each analysis we built a pseudo-population by pooling neurons across patients and selecting a fixed subset of neurons (same subset as relational(cosine) similarity analysis) and concatenating their responses into a data matrix. For every neuron in this subset, trials were grouped by condition; in our case, a condition corresponded to a particular lexical item (for example, the word “man”, “woman”, “king”, “queen”, “boy”, “girl”, “mother”, or “father”), possibly pooled across repeated presentations and surface variants (“man” and “men”). For each neuron and condition we stored a matrix containing all trial-by-trial firing rates. Conditions were organized into matched pairs that instantiate the abstract distinction of interest. A concrete example is a “gender”-like variable defined over four matched word pairs: man–woman, king–queen, boy–girl, and mother–father. In this example, each pair contains a masculine word and a feminine word. More generally, such matched pairs form a dichotomy in our analysis; within each pair one condition is assigned to the “negative” class (for instance the masculine member) and the other condition to the “positive” class (for instance the feminine member). The analysis then asks whether a linear decoder trained to distinguish positive from negative words on some subset of the pairs can correctly classify trails from the left-out pair.
b. Cross-condition generalization with leave-one-pair-out. To quantify CCGP, we used a leave-one-matched-pair-out cross-validation scheme at the level of pairs. Suppose the dichotomy is defined by the four gender word pairs, in one outer fold, we leave out the man–woman pair as the test pair. All trials for “man” and “woman” are held aside, while the remaining three pairs (king–queen, boy–girl, mother–father) are used for training. Within each fold, we enforced equal sampling size across conditions. For every neuron and every condition included in that fold, we randomly subsampled a fixed number of trials per condition without replacement. These subsampled trials were stacked across conditions to form a trial-by-neuron matrix. Trials from all “negative” conditions (e.g. man, king, boy, father) were assigned one label, and trials from all “positive” conditions (woman, queen, girl, mother) were assigned the opposite label. We then standardized each neuron’s activity by z-scoring across trials within that fold. We trained a linear regression with logistic loss on the subsampled training data from the three training pairs and then evaluated it on all subsampled trials from the held-out pair. Accuracy on the held-out pair provides a measure of CCGP for that outer fold. We repeated this procedure, in turn, leaving out each matched pair as the test pair (so we also evaluated generalization to king–queen when the other pairs were used for training, and so on) and averaged accuracy across these outer folds to obtain one CCGP value for that bootstrap iteration. To obtain a stable estimate that is not driven by a particular choice of trials, we repeated the entire subsampling and left-one-pair-out procedure 600 times, each time drawing a new random subsample of trials per condition. The primary CCGP estimate reported for each population and dichotomy is the mean accuracy across these iterations.
c. Regularized linear decoding and nested cross-validation. To avoid overfitting idiosyncratic fluctuations in the training data, decoding was performed with a linear classifier with logistic loss and L2 regularization. However, the optimal strength of regularization is not known in advance and may differ across populations and dichotomies. To choose the regularization strength in a data-driven way, we used nested cross-validation inside the outer leave-one-pair-out loop. Returning to the four-pair example, consider again the outer fold where man–woman is held out as the test pair. The remaining three pairs—king–queen, boy–girl, and mother–father—serve as the training pairs. Within this training set, we perform an inner leave-one-pair-out cross-validation to select the regularization parameter. For a candidate value of λ, first, we hold out one of the training pairs as an inner validation pair, for example boy–girl. We train the classifier on subsampled and standardized trials from the remaining pairs (king–queen and mother–father) using that λ, then measure how well it predicts the labels for the held-out validation pair boy–girl. We repeat this inner procedure holding out each training pair in turn (boy–girl, king–queen, mother–father), and average the validation errors across these inner folds to obtain a cross-validated error for that λ. We repeat the inner leave-one-pair-out procedure for a small grid of candidate λ values (1/N, 3e−3,1e−2, 3e−2, 6e−2, 1e−1, 3e-1, where N is the number of neurons) The λ that yields the lowest average validation error is selected for that outer fold; if multiple λ values perform equally well within numerical precision, we choose a value near the center of this set on a logarithmic scale.
d. Geometric (neuron-shuffle) null. To determine whether high CCGP truly reflects meaningful structure in the population code, we compared the empirical CCGP to a geometric null model that disrupts the consistent identity of neurons across conditions while preserving activity distributions within each condition. For this null, we repeated the entire analysis described above, but before decoding we shuffled neuron identities within each condition. Concretely, for a given condition (e.g. all trials of “man”), we assembled a trial-by-neuron matrix and randomly permuted the order of neurons (columns) within that matrix. This operation was performed independently in each condition (man, woman, king, queen, etc.). As a result, each neuron’s pattern of responses is effectively reassigned to a different label in each condition, preserving the distribution of firing rates across trials and across neurons within each condition but destroying the consistent mapping between neurons and their tuning across different words and pairs. After this neuron shuffling, we ran the same nested leave-one-pair-out decoding procedure as for the empirical data, including the same trial subsampling and regularization selection. The resulting distribution of CCGP values across iterations provides a geometric null that captures how much cross-pair generalization would be expected from high-dimensional variability alone, in the absence of stable neuron-specific codes that align across conditions.
e. Label-shuffle null. As a complementary control, we also constructed a label-shuffle null model that preserves the neural data but destroys the relationship between neural activity and the semantic variable being decoded. In this variant we used the original, unshuffled neural responses but, within each outer fold, randomly permuted the class labels assigned to trials before training and testing the decoder. The entire nested cross-validation procedure was then run on these randomly relabeled data. This null provides a baseline for CCGP that would be expected if the decoder were applied to the same neural activity but the mapping between population patterns and the conceptual dichotomy (e.g. masculine vs feminine words) were arbitrary.
f. Summary and statistical comparison. For each analogy, we therefore obtained three CCGP distributions across bootstrap iterations: one from the intact data, one from the neuron-shuffle (geometric) null, and one from the label-shuffle null. The empirical CCGP for a given population was taken as the mean accuracy across iterations on the intact data. To assess significance, we compared this value to the corresponding null distributions by computing the proportion of null iterations whose CCGP was at least as large as the empirical mean, with a small finite-sample correction. In the man–woman, king–queen, boy–girl, mother–father example, a high and significant CCGP means that a linear decoder trained to distinguish three of these word pairs reliably predicts the left-out pair, in a way that cannot be explained by shuffled labels or by random reassignments of neuron identity.

### Additive factor model of pronoun-evoked population activity

We quantified the extent to which pronoun-evoked hippocampal population activity could be explained by an additive (factorized) representation of grammatical features versus feature interactions. The analysis was performed at the individual-trial level using the trial-by-neuron firing-rate matrix 𝑍 (restricted to 12 pronoun categories).Let 𝑇 be the number of retained trials and 𝑁 the number of neurons. The firing-rate matrix is

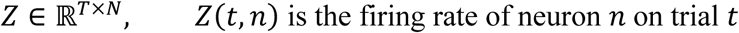

Trials were restricted to the 12 pronoun categories

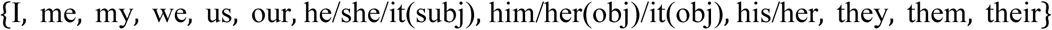

Each retained trial 𝑡 was assigned three categorical grammatical factors based on pronoun identity. Case

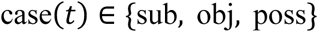

with pronoun groupings

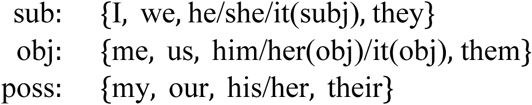

Number

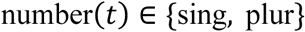

with pronoun groupings

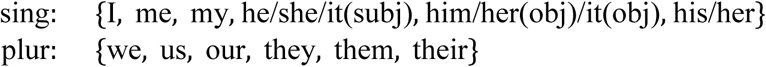

Person

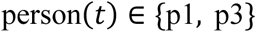

with pronoun groupings

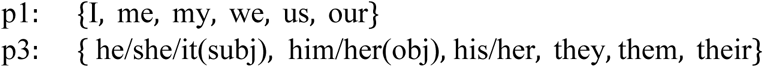

We fit multivariate linear regressions predicting the 𝑁-dimensional population response on each trial from dummy-coded grammatical factors. All models were fit simultaneously across neurons by ordinary least squares (OLS), using the same design matrix for all neurons.The main-effects model included an intercept plus additive terms for case, number, and person. Using reference coding with baselines

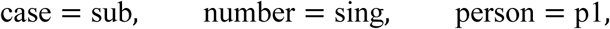

the main-effects design matrix contained the regressors

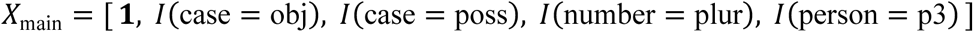

where 𝐼(⋅) is the indicator function. Let 𝑋_main_ ∈ ℝ^T×*p*^_main_ and 𝐵_main_ ∈ ℝ*^p^*^main×*N*^ be the coefficient matrix. Coefficients and predictions were computed by OLS (MATLAB mldivide notation):

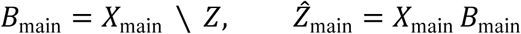

The full model expanded the design matrix with interaction terms up to three-way. The additional regressors were:

**case** × **number**

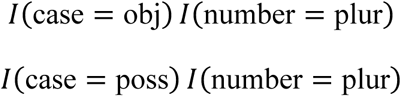

**case** × **person**

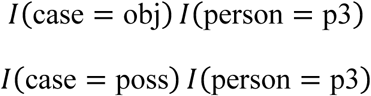

**number** × **person**

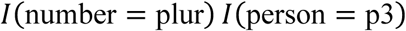

**case** × **number** × **person**

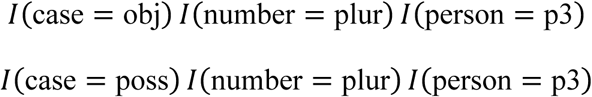

Let 𝑋_full_ ∈ ℝ^*T*×*P*full^ be the resulting design matrix. Coefficients and predictions were computed analogously:

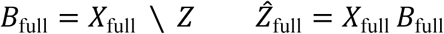

Model fit was summarized as a single population-level 𝑅^2^by pooling squared errors across all trials and neurons. For a generic prediction 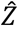, we computed

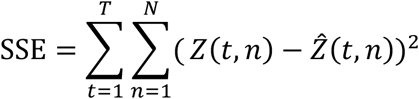

and

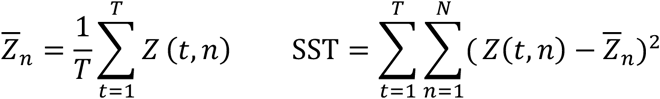

The pooled coefficient of determination was then

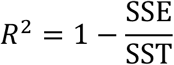

We computed 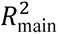 using 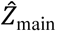 and 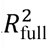 using 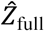 Because the full model includes additional degrees of freedom, 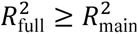. To quantify the relative contribution of additive structure within the explainable component, we also computed

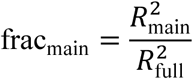

Statistical significance of additive factorization was assessed using a label-shuffle null distribution.

On each permutation, we randomly permuted the trial-wise word labels label(t) across the retained trials (keeping Z fixed), recomputed case(t), number(t), and person(t) from the shuffled labels, rebuilt 𝑋_main_ and 𝑋_full_, refit both models by OLS, and recomputed 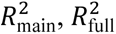, and frac_main_. This procedure preserves the marginal distribution of firing rates and trial counts while destroying the correspondence between population activity and pronoun identity/feature structure The observed frac_main_ was compared to the permutation distribution using a one-sided test (higher frac_main_ indicates a more factorized, main-effects–dominant structure). With 𝑁_perm_ shuffles, the p-value was computed as:

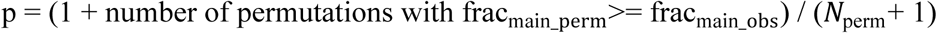

### Differences across brain areas

We tested whether the prevalence of significant word-pair effects differed between hippocampus (HPC) and anterior cingulate cortex (ACC) across 15 analogy types. Same sets of analysis was repeated comparing HPC to OFC. To make the regional comparison robust to differences in neuronal sample size, we summarized each word pair with a 0–1 significance-prevalence score that reflects how likely that pair is to be deemed significantly aligned with other pairs in an analogy under ACC neuron count-matched sampling. Concretely, this score corresponds to whether a pair is significant in ACC, and—because HPC had a larger neuronal pool—to the fraction of neuron-matched resamples (500 resamples) in HPC that yield a significant effect. This yields a common metric across areas in which larger values indicate more reliably detectable word-pair effects at comparable sampling, rather than simply reflecting differences in the number of recorded neurons.

We fit a linear mixed-effects model with fixed effects of Area, Type, and their interaction, and a random intercept for word-pair identity (pairID) : sigPrev∼1+Area×Type+(1∣PairID), 412-word pair observations; Residual method for the degrees of freedom. Marginal ANOVA tests indicated significant effects of Area, Type and a robust Area × Type.

To complement the linear mixed-effects analysis, we performed a brain area label-shuffling, size-matched resampling test that asks whether HPC and ACC show different *profiles* of significant word-pair counts across the 15 analogy types, independent of unequal neuron counts. For each area we computed a 15-element vector, vArea=[sigCount1,…,sigCount15] where each element is the number of significant word pairs (*sigCount*) for that analogy type. We quantified the similarity between the two area profiles using cosine similarity between the HPC and ACC vectors (lower cosine similarity = more different profiles).

Because the number of recorded neurons differed between areas, we used balanced resampling: in each iteration we randomly downsampled HPC neurons to match the ACC neuron count, recomputed the HPC and ACC *sigCount* vectors, and then computed cosine similarity. We repeated this procedure 500 times and used the average cosine similarity across the 500 downsamplings as the observed (non-shuffled) profile similarity.

To generate a null distribution under the hypothesis that area identity does not matter (i.e., neurons are exchangeable between HPC and ACC), we pooled neurons across areas, randomly permuted (shuffled) the area label assigned to each neuron, then repeated the same size-matched sampling and cosine-similarity computation 500 times. We then asked how often the shuffled data produced a profile similarity as low as (or lower than) the observed value—i.e., a pattern as different or more different than the real HPC vs ACC profiles.

**Supplementary Figure 1.**
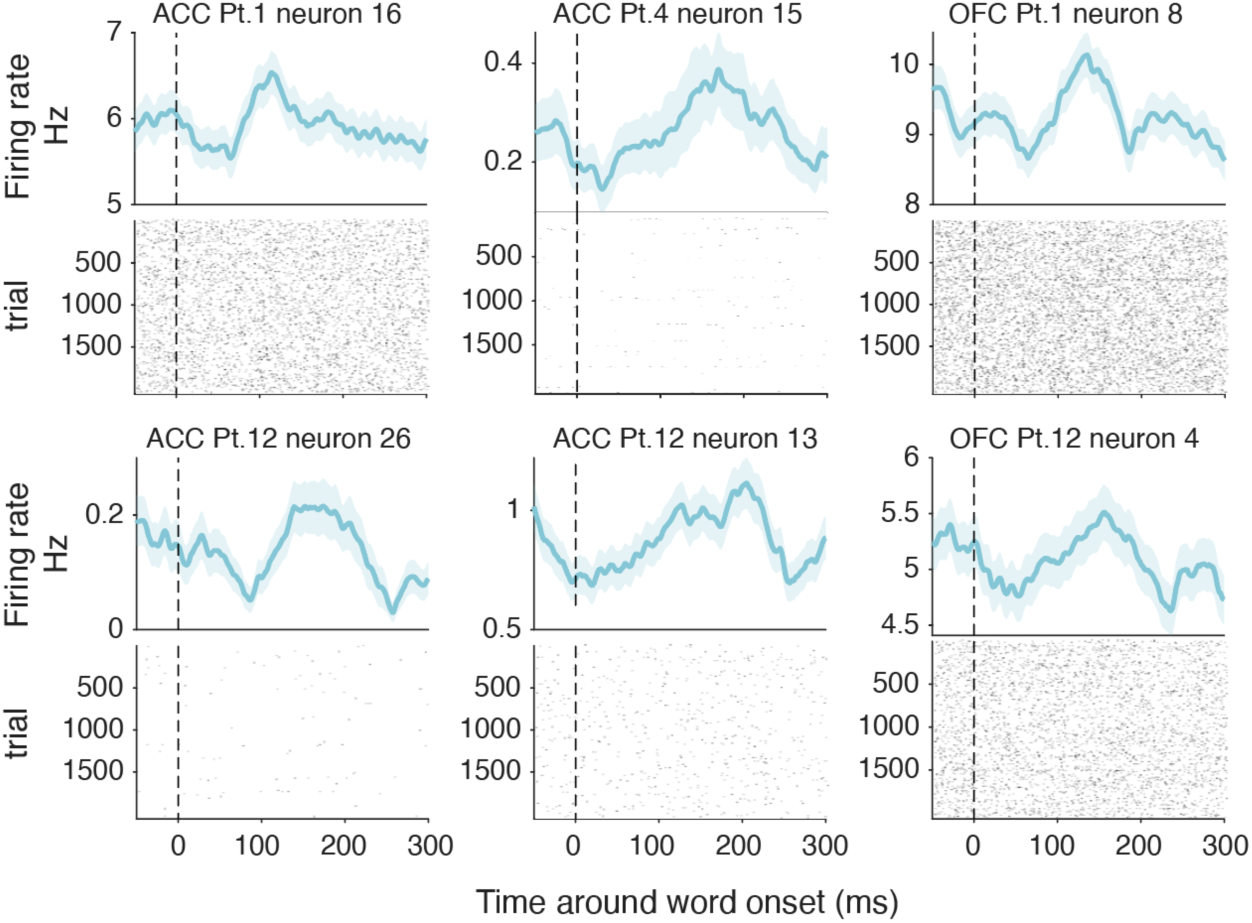
| PSTH for example neurons in other brain areas. Same as Figure 1C but for example ACC/OFC neurons.

**Supplementary Figure 2.**
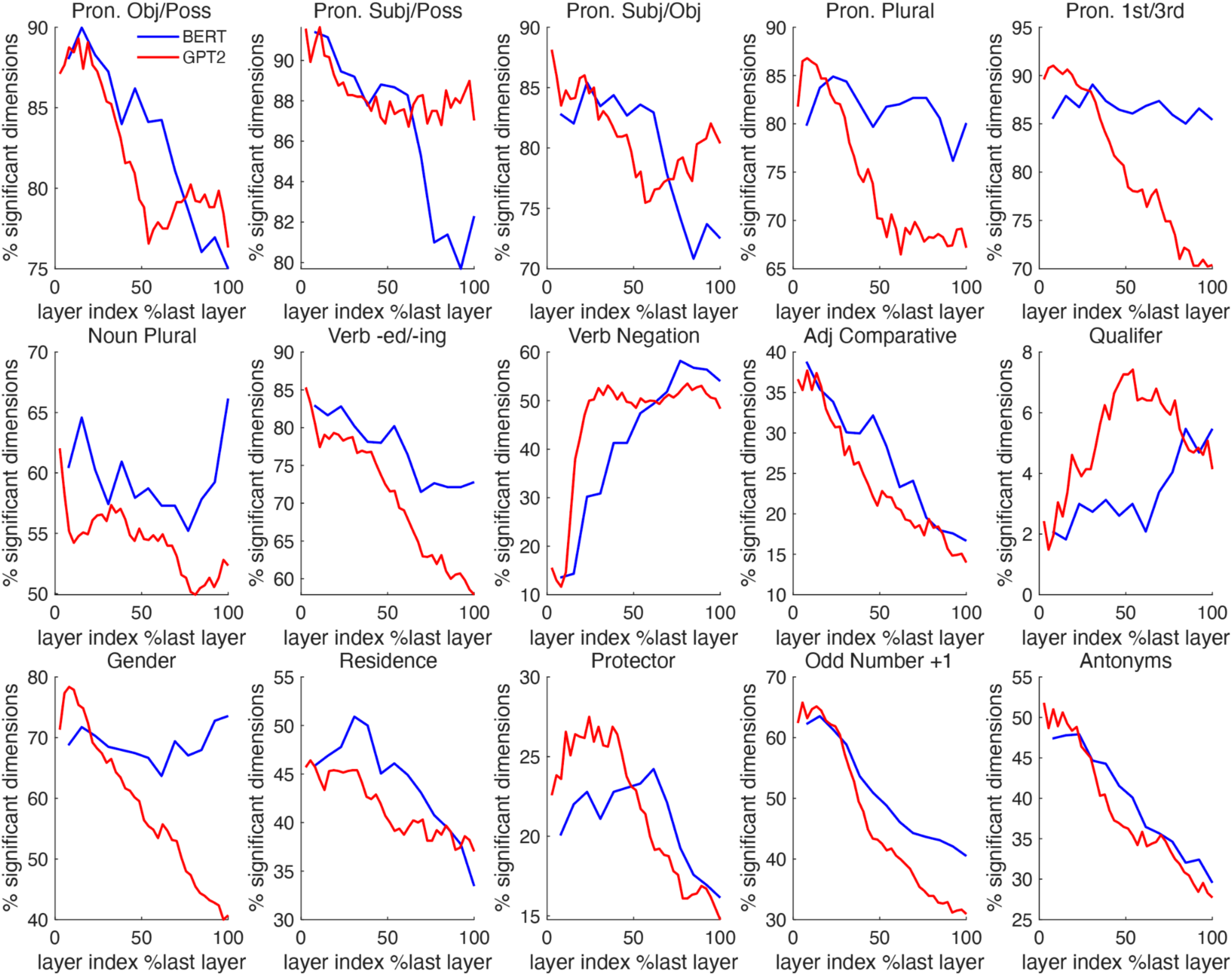
| Analogical representation across BERT and GPT layers. We analyzed 13 layers of BERT (blue) and 37 layers of GPT-2 (red), normalizing the x-axis by relative depth (e.g., the 6th layer of BERT appears at 0.46). Across 15 analogy types, the fraction of dimensions meeting our criterion generally dropped as layers deepened (increased layer index). This suggests that later layers rely on fewer specialized embedding dimensions. Exceptions include context-dependent operations (e.g., negation (“feel” vs “(not) feel” and qualifier “fall” vs “(kind of) fall” modification), where performance depends on contextual binding.

**Supplementary Figure 3.**
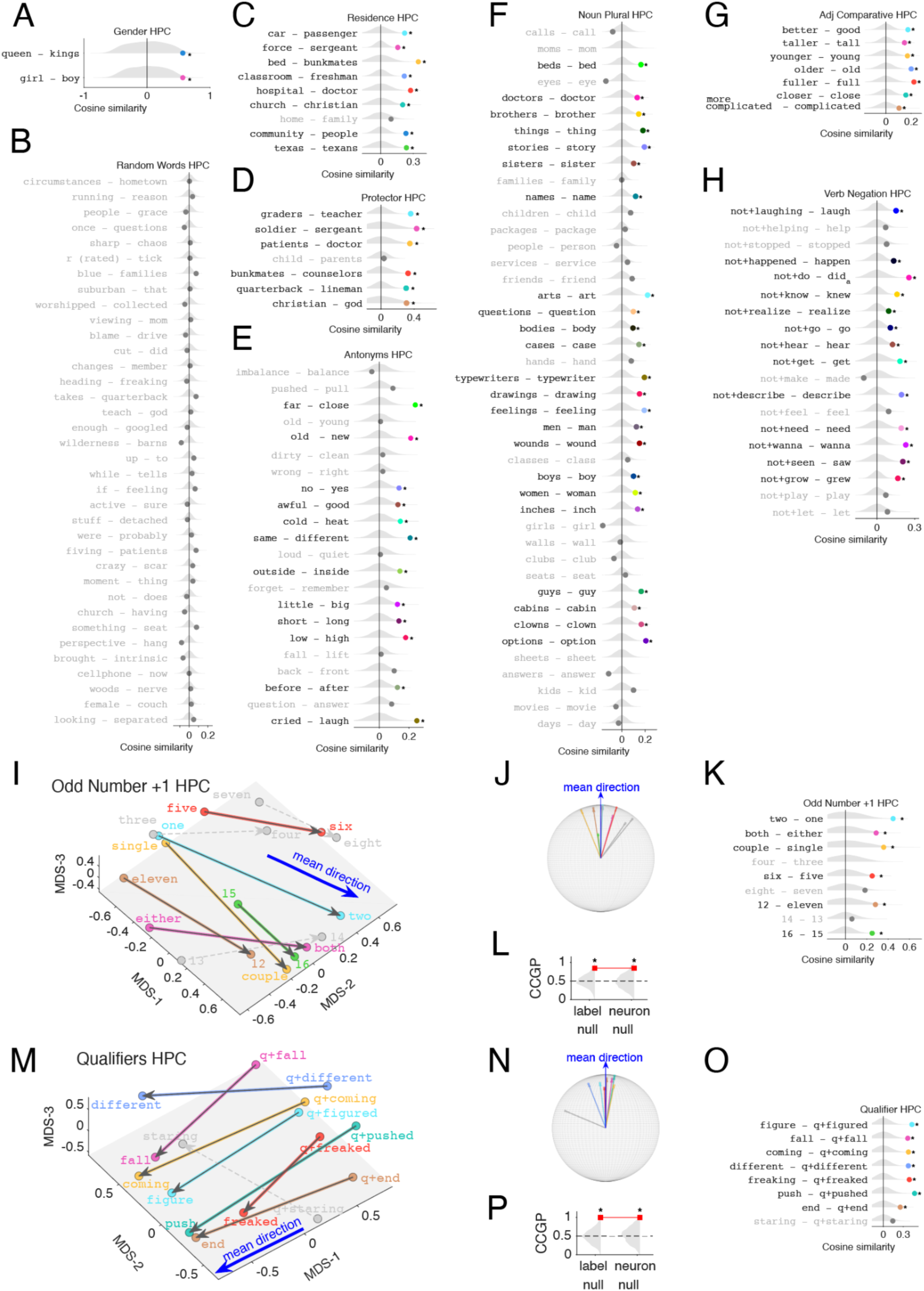
| Additional hippocampal analogy cosine similarities/additional analogy types supporting Fig2 and Fig3 **A–H, Alignment of individual word-pair difference vectors to the other word pairs within-category in HPC.** For each semantic relationship, we computed leave-one-out mean difference directions from high-dimensional neural word embeddings (population firing-rate vectors) and measured mean cosine similarity between each held-out word-pair difference vector and the training directions. Grey violins indicate a random word-pair null distribution; dots indicate observed similarities (colored: significant after Benjamini–Hochberg FDR correction across pairs within category; grey: non-significant). **A,** Gender (subset shown). **B,** Random word pairs (control), only 35 pairs shown, see Figure 2 for the rest 15 pairs. **C–H,** Residence, protector, antonyms (valence-oriented), noun plural, adjective comparative, and verb negation. These panels correspond to the alignment analyses shown in the main figures; see Figure 3 for the accompanying MDS and CCGP for these relationships. **I–P, Two additional semantic relationships show consistent axes and support cross-condition generalization.** We identified aligned difference directions for **odd number + 1** and for **qualifiers** (words preceded by *kind of* / *slightly*; shown as “q+” prefixes). **I,M,** 3D MDS visualization of neural word embeddings with arrows indicating paired difference vectors (MDS shown for visualization only; grey planes are visual aids). Colored arrows/labels denote pairs whose high-dimensional difference vectors are significantly aligned to the within-category directions (permutation test against random word-pair null; Benjamini–Hochberg FDR across pairs within category); grey arrows/labels denote non-significant pairs. Blue arrows indicate the mean direction projected into the MDS view. **J,N,** The same vectors after tail alignment and rigid rotation so the mean direction points upward, with vector lengths normalized to a unit sphere (visualization only; angles preserved). **K,O,** mean Cosine similarity of each pair to the leave-pair-out directions from other word pairs in the same category with null distributions (grey violins). **L,P,** Cross-condition generalization performance (CCGP) for each relationship (red marker), compared to label-shuffle and neuron-shuffle nulls (grey violins; dashed line indicates chance). Asterisks indicate p < 0.05 (permutation test; n shuffles = 600) relative to the corresponding null.

**Supplementary Figure 4.**
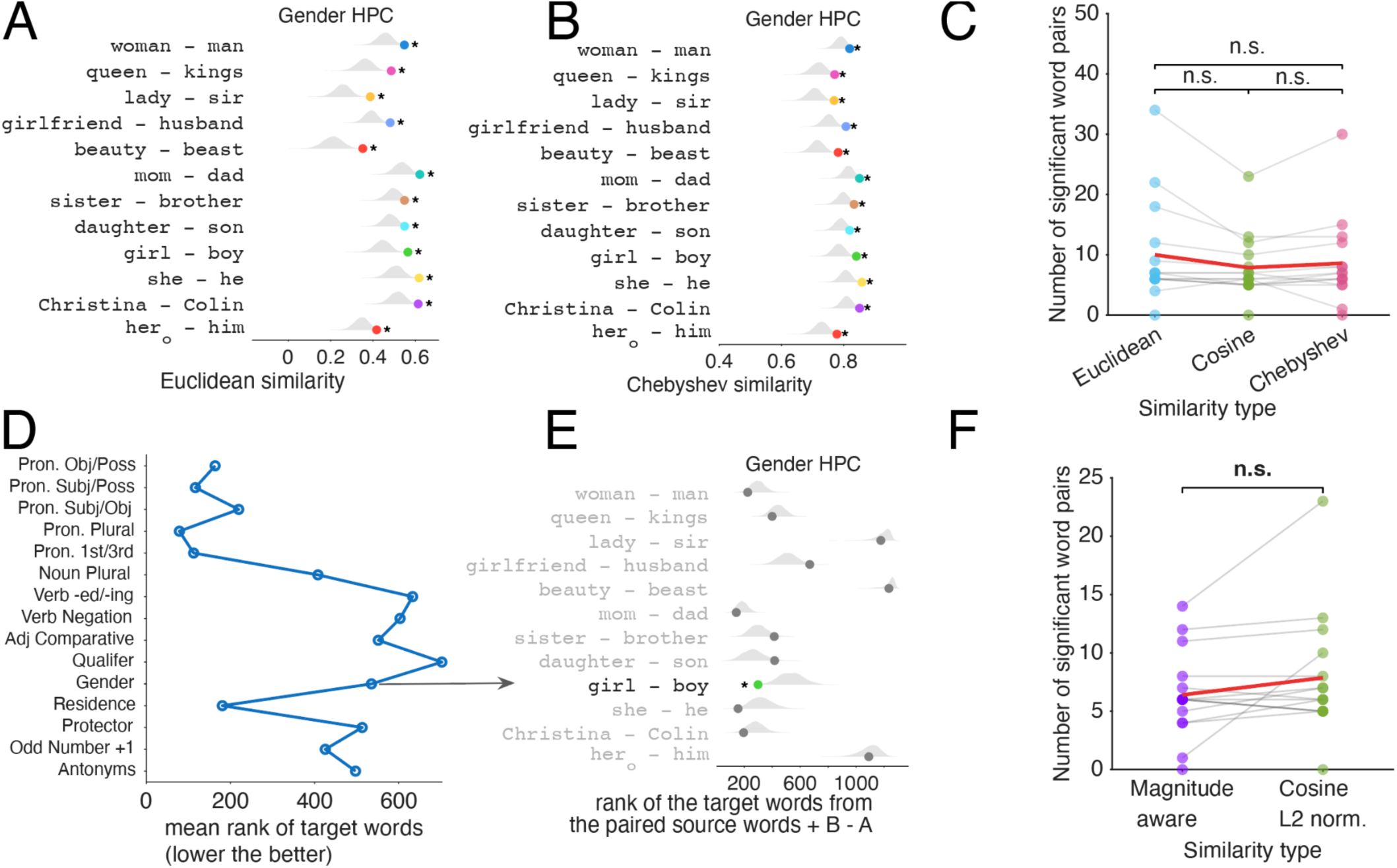
| Robustness to similarity metric and limited analogy retrieval from neural vector arithmetic. **A–C, Distance-metric robustness for alignment.** For the gender analogy set in hippocampus (HPC), pairwise alignment between within-category difference vectors was recomputed using alternative similarity metrics. **A,** Euclidean similarity (1 - euclidean distance) between each gender difference vector (e.g., “*woman*”−“*man*”) and the mean leave-one-out gender vectors; grey violins indicate a random-pair null distribution and colored dots show observed similarity (asterisks: significant after within-category multiple-comparisons control, as in the main analysis). **B,** Same analysis using 1 - Chebyshev distance. Across analogy types, the number of significant aligned word pairs is comparable under Euclidean, cosine, and Chebyshev similarity (paired Wilcoxon signed-rank tests, n.s.). **D–F, Analogy retrieval is weak and insensitive to scoring choice. D,** For each analogy type, mean rank of the correct target word under vector-arithmetic retrieval: for each pair A:B and source word C, the model predicts a target D from D(estimate)=(B−A)+C and ranks all candidate words in the corpus by similarity to D(estimate) (lower rank indicates better retrieval). **E,** Example retrieval rank distributions for the gender set; only “*girl*”–“*boy*” is retrieved significantly above chance relative to a random-difference baseline. **F,** The number of significant retrieved word pairs is not significantly different when using a magnitude-aware scoring rule versus cosine similarity after L2 normalization (paired Wilcoxon signed-rank test, n.s.). Supplementary result and discussion for this figure: We primarily relied on cosine similarity to quantify vector-offset analogies, as it remains the standard in the field (Mikolov et al., 2013). However, given that this metric could be sensitive to normalization and scoring choices (Levy & Goldberg, 2014), we investigated whether our neural axis-alignment findings were robust across metrics or merely artifacts of specific distance calculations. Recomputing pairwise alignment with Euclidean and Chebyshev similarity (1 - distance) produced the same qualitative outcome and did not significantly change the number of significant aligned pairs across analogy types (**Supplementary Figure 4**, Wilcoxon signed-rank tests, p > 0.05). We also evaluated a stricter “analogy retrieval” criterion—ranking the true target word against all other words using vector arithmetic (B − A + C)—and found that retrieval was only modestly above chance: across analogy families, the target routinely ranked well outside the top set of candidates (typically >100), and in the gender set only “*girl*”–“*boy*” was reliably retrieved above chance (**Supplementary Figure 4**). The number of significant word pairs did not alter by switching from cosine similarity scoring to this analogy retrieval regime, reinforcing that our primary alignment results are largely distance-metric or paradigm agnostic, while leaving open what similarity computation downstream circuits implement and whether the exact parallelogram-style retrieval is the algorithm the brain uses to solve SAT style analogy reasoning tasks.

**Supplementary Figure 5.**
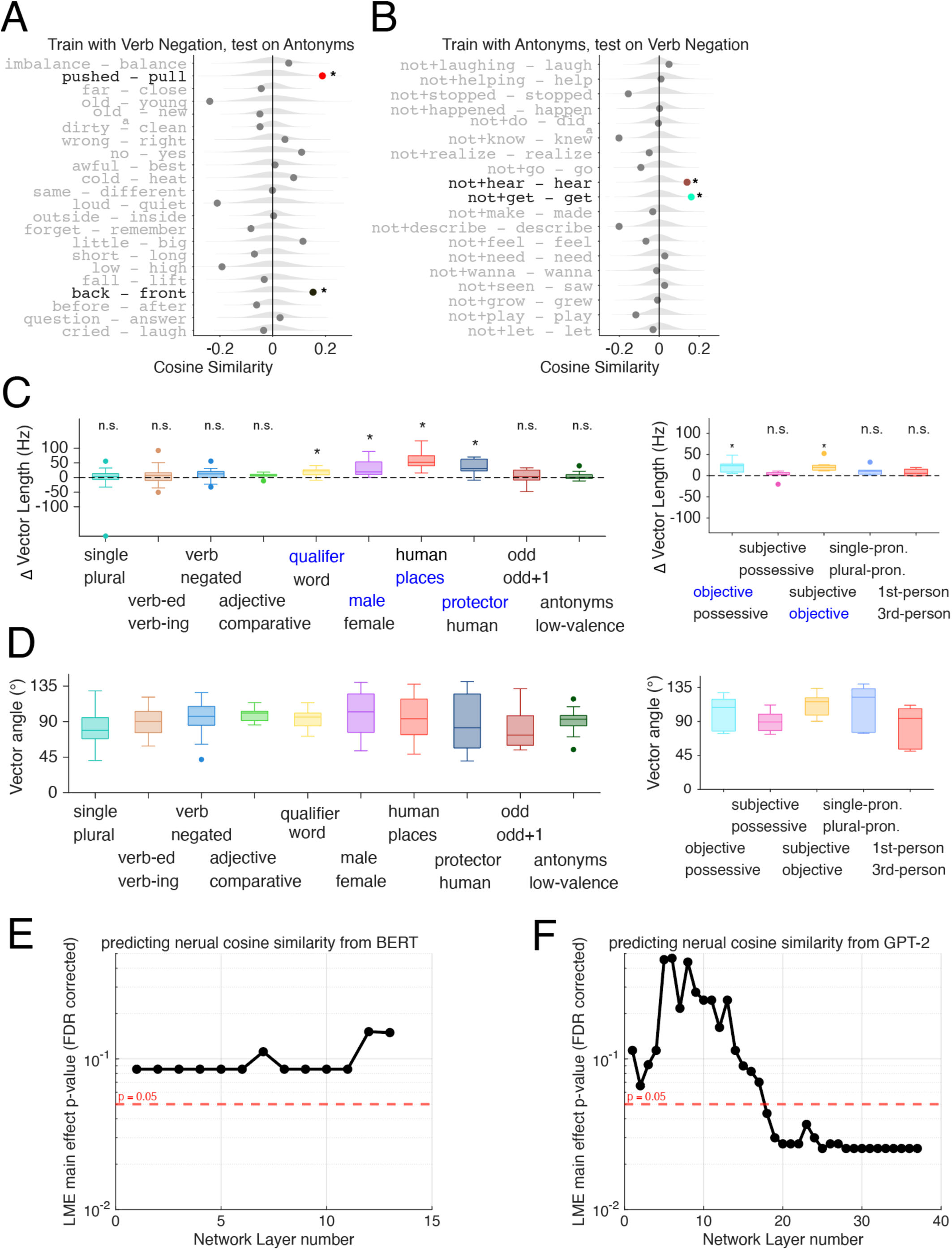
| Distinct semantic axes and control analyses for vector geometry, with additional pronoun results supporting **Figure 2-4** **A–B, Antonym and verb-negation axes do not generalize to one another.** We tested whether the coding directions learned from one analogy family predicts pairwise directions in another by training the directions on one set of word-pair difference vectors and then computing cosine similarity between those directions and each held-out pair in the other set (grey violins: random-pair null; dots: observed mean similarities; asterisks: significant alignment vs. null after multiple-comparison correction within panel). **C–D, Axis alignment is not explained by systematic changes in vector magnitude or simple sign inversions. C,** Signed differences in raw vector length (Δ vector length; units reflect firing-rate vector magnitude) between the two words within each pair, computed consistently using the top 30 tuned (lowest p(AUC)) neurons for that analogy type. Blue labels indicate which side of the pair had the larger mean length; asterisks denote analogy types with a significant nonzero mean Δ length (paired t-test across word pairs, Benjamini–Hochberg FDR). For visualization only, pronoun analogies are plotted separately from the other analogy types (the repeated y-axis is identical). Because our main analyses L2-normalize each word vector before subtraction (**Methods**), systematic differences in raw vector length cannot account for the observed axis alignment. **D,** Angles between the two raw word vectors within each pair. Negation (e.g., “not *grow*” vs. “*grow*”) and comparative morphology (e.g., “*taller*” vs. “*tall*”) are not consistent with antipodal inversions (∼180°), but instead tend to be approximately orthogonal (∼90°) to the corresponding root-word vectors (cf. Zuanazzi et al., 2024). **E-F,** Per layer linear mixed effects model on HPC cosine similarities of individual word pairs across all fifteen diverse analogies. The model was specified as HPC ∼ 1 + LLM + (LLM | analogy type), where we assessed the fixed effect of the LLM similarity while accounting for random intercepts and slopes across analogy types, reporting the FDR corrected p-value of the main effect for each layer of BERT (**E**) or GPT-2 (**F**).

**Supplementary Figure 6.**
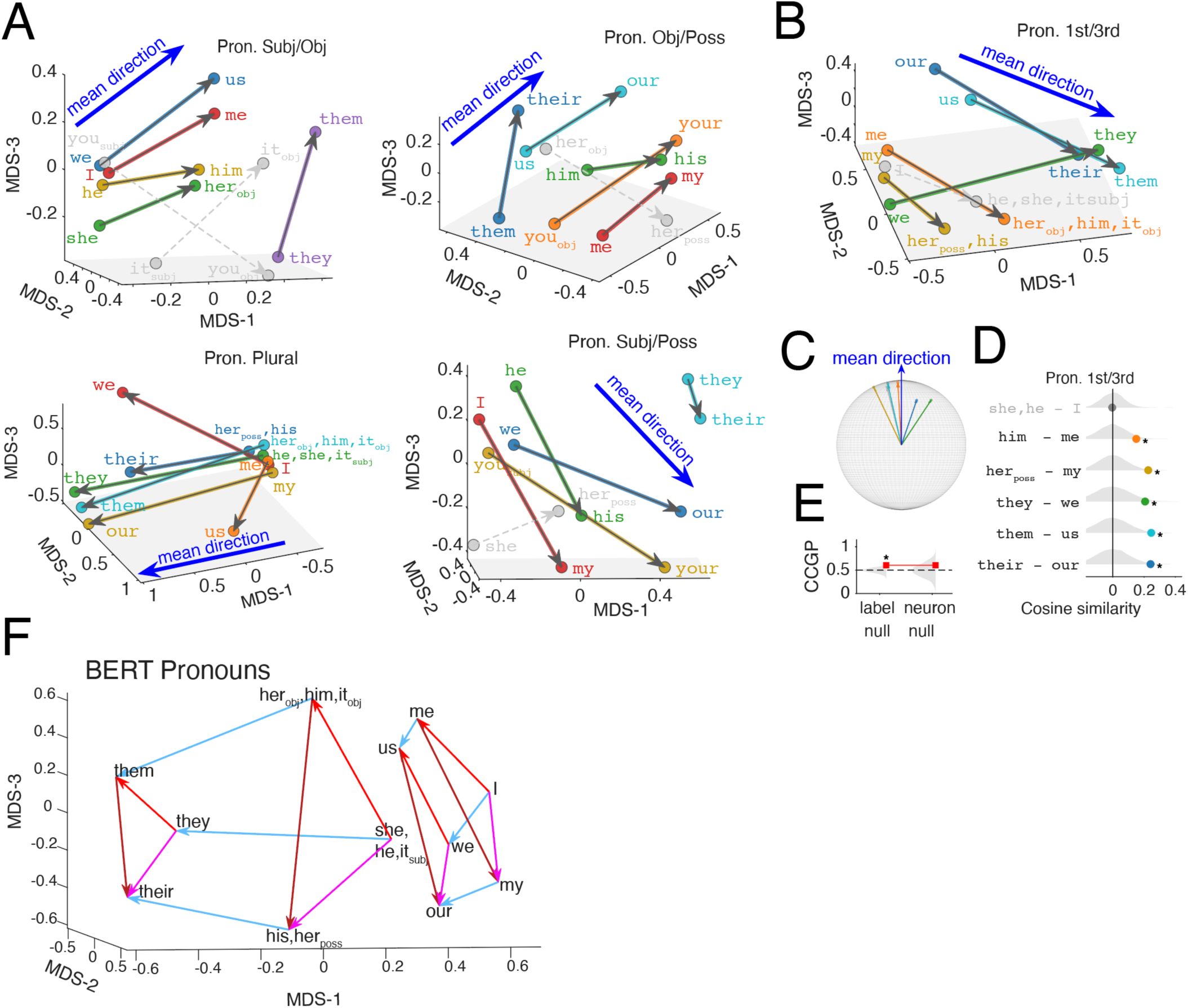
| Semantic axes of pronouns in an LLM, supporting **Figure 4** **A, Pronoun analogies shown in MDS for visualization.** Low-dimensional MDS plots for four pronoun analogy families (subject/object, object/possessive, plural, and subject/possessive). These MDS visualizations are provided for reference; associated mean-direction, cosine-alignment, and CCGP analyses are shown in Fig. 4. **B–E, First-/third-person pronoun analogy. B,** MDS visualization for the first-/third-person axis. **C,** Tail-aligned difference vectors projected onto a unit sphere and rotated so the mean direction points upward. **D,** mean Cosine similarity of each first-/third-person pair to the leave-one-out training directions (rest of the word-pairs) (null distributions in grey); 5/6 pairs are significantly aligned (binomial test p < 0.001). **E,** Cross-condition generalization performance (CCGP) for first vs. third person, shown relative to label-shuffled and neuron-shuffled null distributions. Asterisks indicate p < 0.05 (permutation test; n shuffles = 600) relative to the corresponding null. **F**, Same analysis of Figure 4 **M-O** but in BERT. MDS visualization of BERT contextual embeddings for the corresponding pronoun set. n = 52 BERT units (top 30 most tuned units from each of the 4 analogies in panel Figure 4A-L pooled). Arrows use the same color conventions as in Figure 4 **M-O**, and the embedding exhibits a similar prism-like organization with more clearly separated first- and third-person configurations.

**Supplementary Figure 7.**
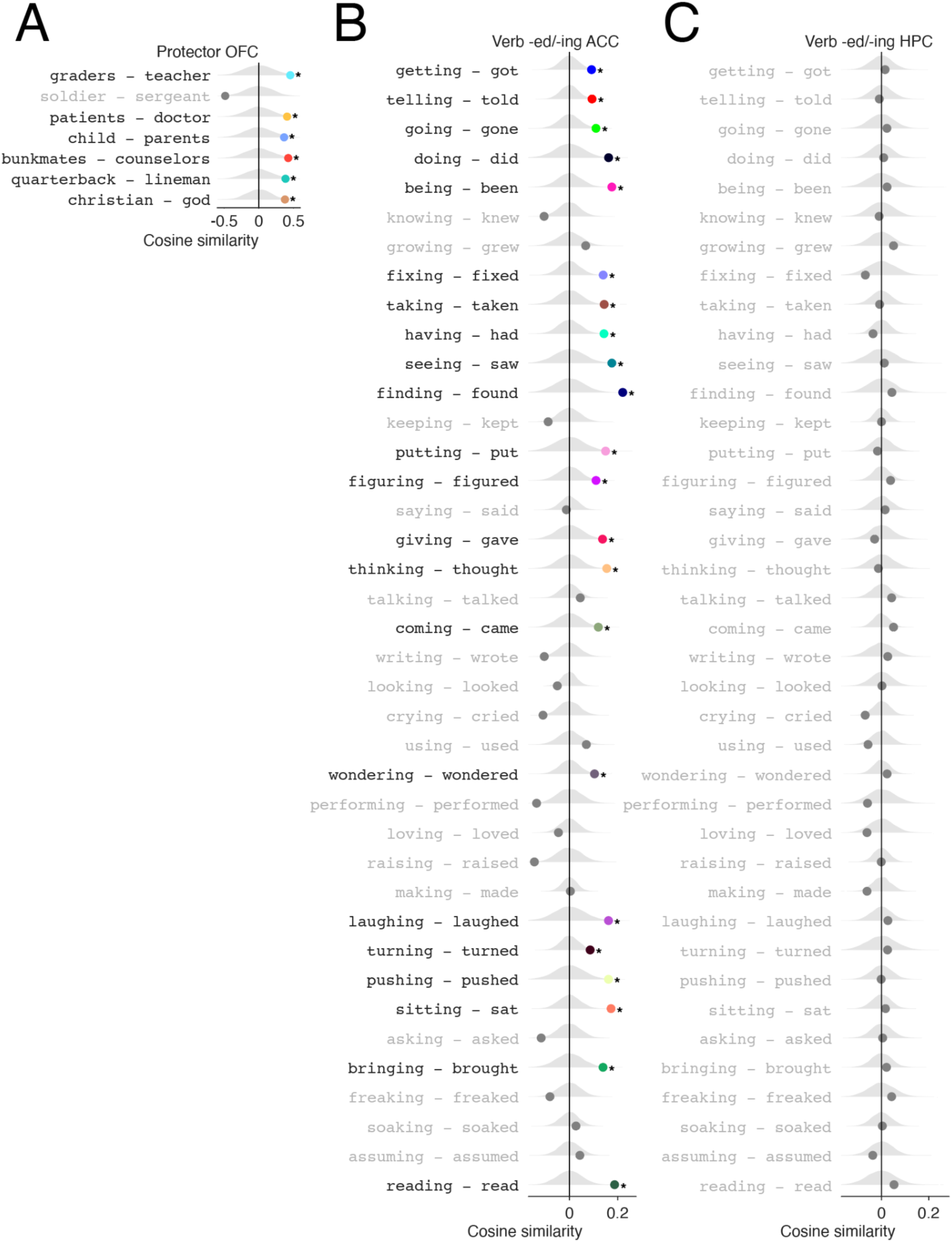
| Pairwise cosine-similarity evidence for analogy axes in ACC and OFC supporting **Figure 5**. **A,** Protector analogy in orbitofrontal cortex (OFC). For each protector word pair, we computed the mean cosine similarity between that pair’s high-dimensional population difference vector and that of remaining word-pairs (leave-one-out) within OFC. Grey violins show a null distribution obtained from random word-pair difference vectors; dots indicate observed similarities. Colored dots/labels mark pairs that are significantly aligned with the rest of the pairs after Benjamini–Hochberg FDR correction across pairs within the category; grey labels indicate non-significant pairs. Dot colors match the corresponding vectors/labels in the MDS visualization in Figure 5. **B–C,** Verb *-ed*/*-ing* analogy in anterior cingulate cortex (ACC) and hippocampus (HPC). Same analysis as in **A**, shown for the verb *-ed*/*-ing* relationship in **ACC** (**B**) and **HPC** (**C**). In ACC, many verb pairs exhibit significant alignment to the direction of other *-ed*/*-ing* word pairs (colored points), whereas in HPC the corresponding cosine similarities cluster near the null distribution and do not show robust pairwise alignment. Colors again correspond to the matched word pairs/vectors in the MDS panels in Figure 5.

**Supplementary Figure 8.**
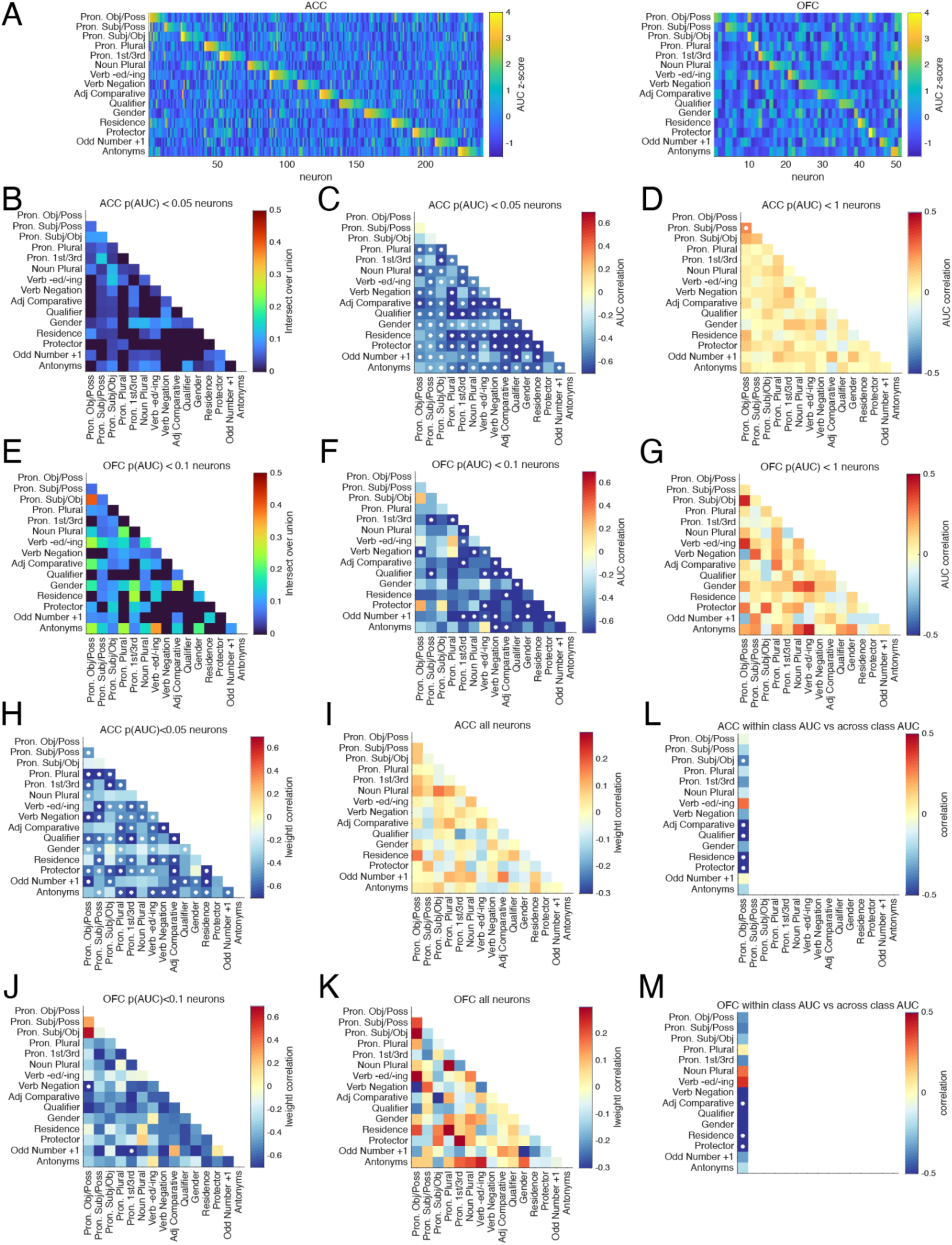
| Specialization in OFC and ACC neurons. Same specialization results as in Figure 6 but for OFC and ACC neurons.

**Supplementary Table 1.**
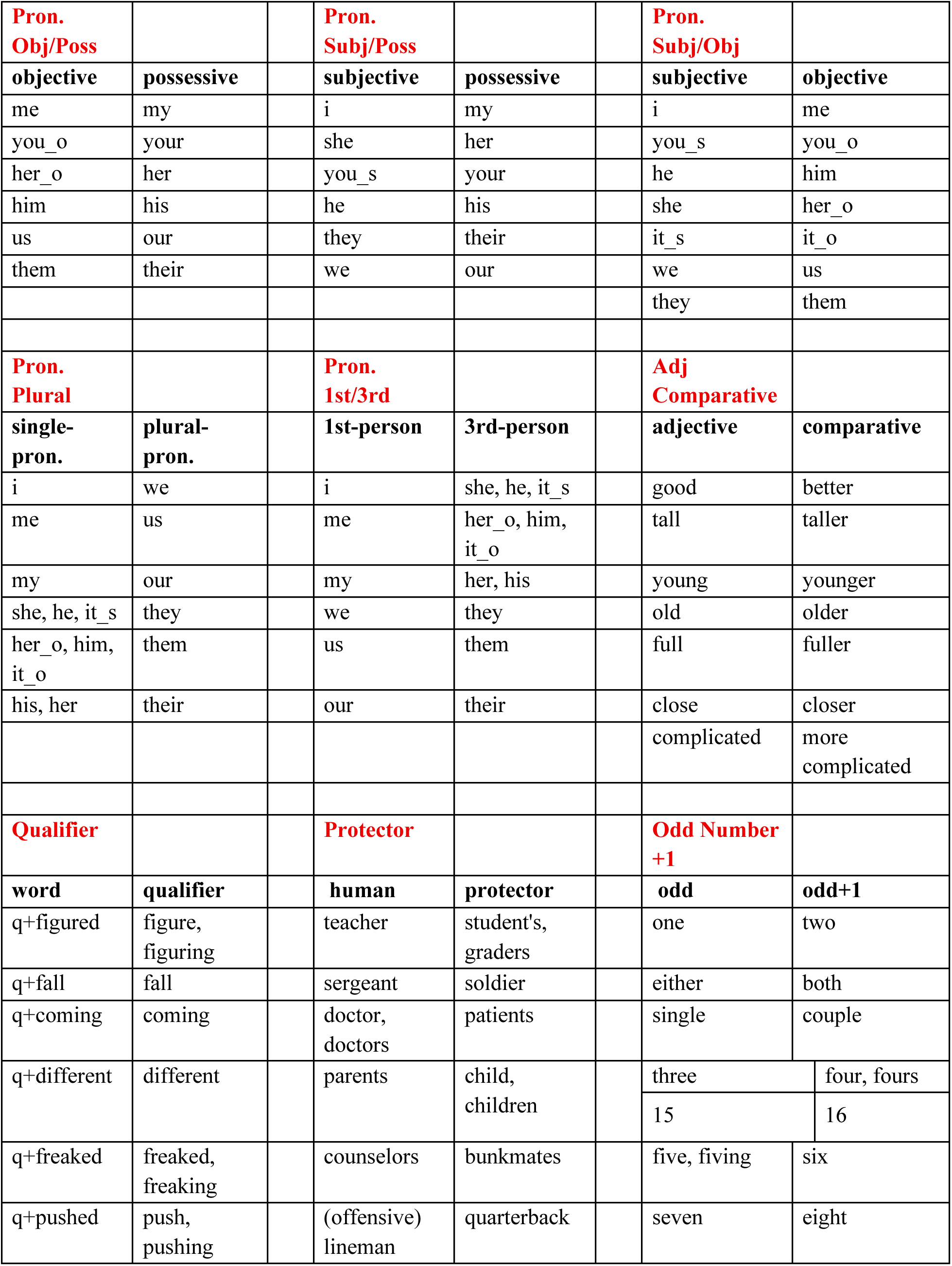
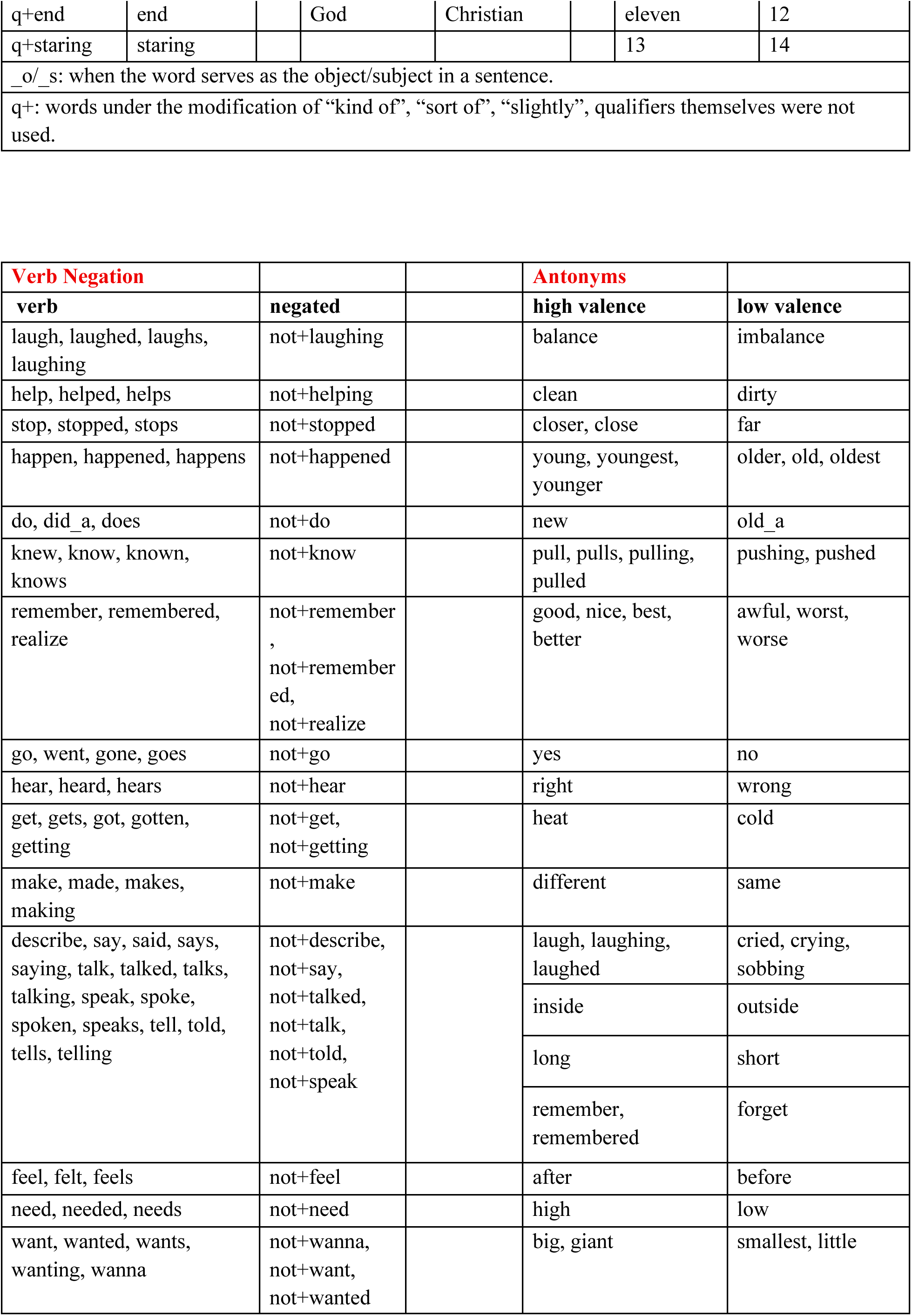
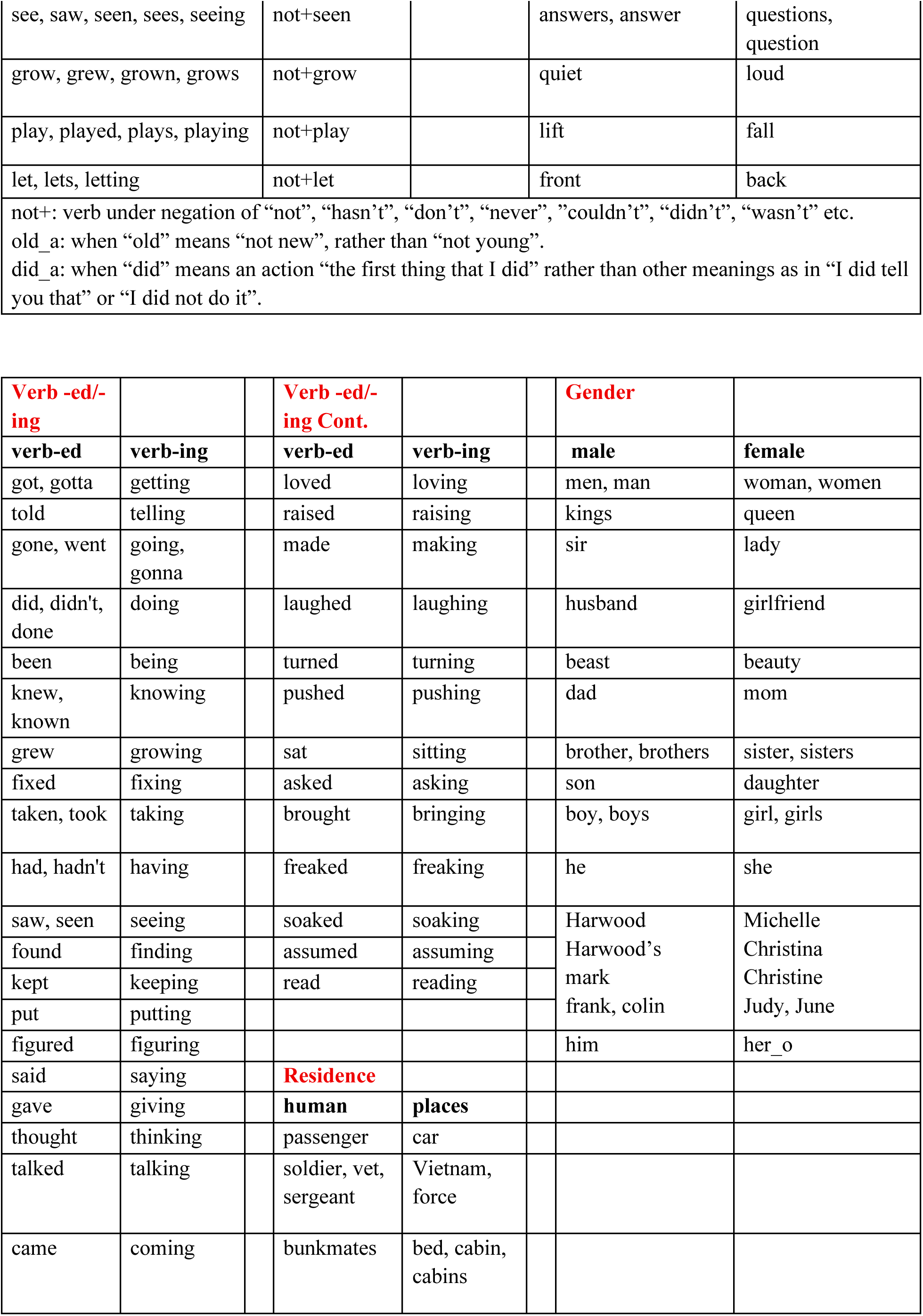
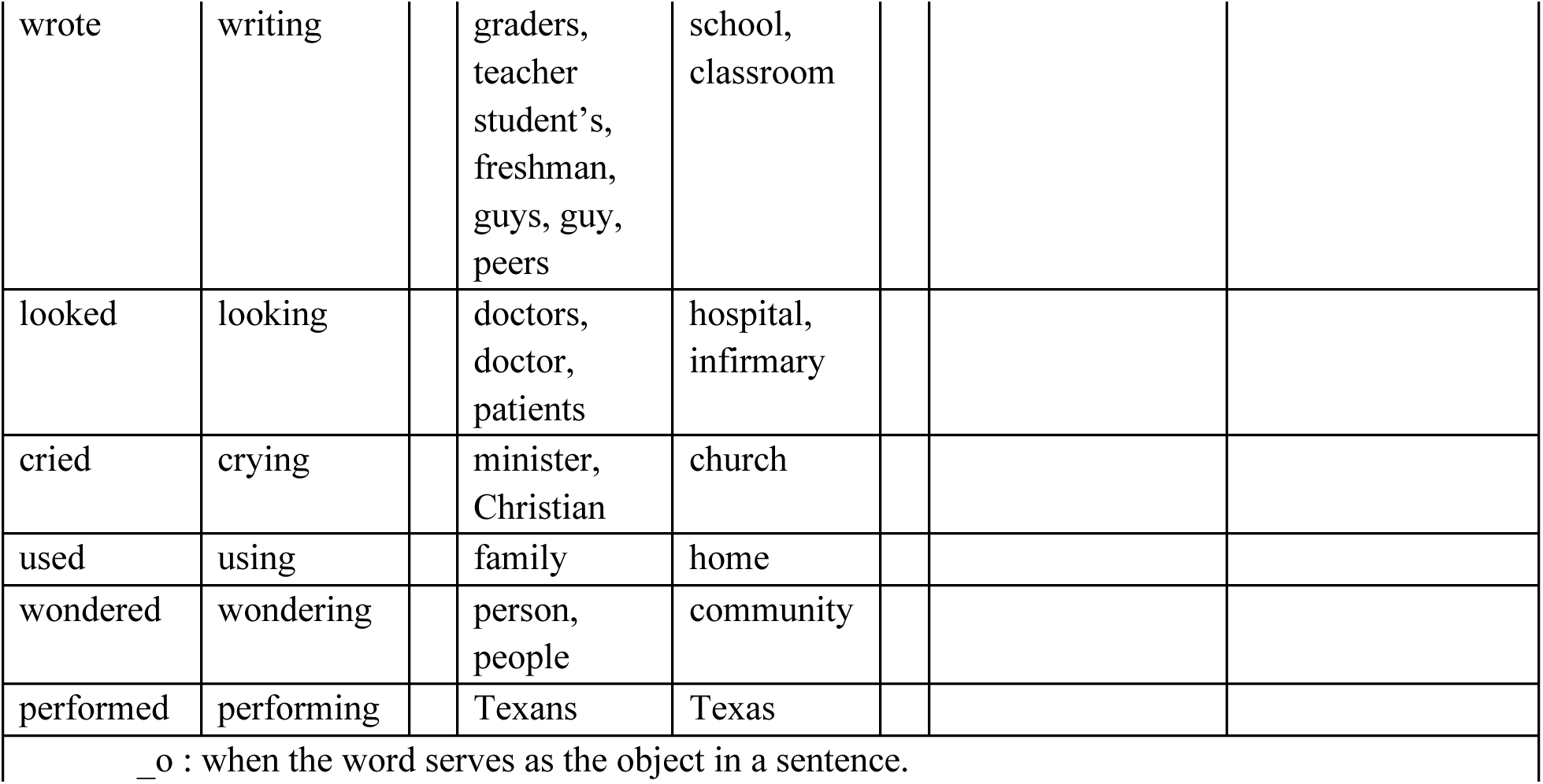
Full list of words used for 14 analogies. see Supplementary Figure 3F for words used in noun single plural analogy.

